# Cycles upon cycles - Temperature Scaling of Medaka Development

**DOI:** 10.64898/2026.04.29.721546

**Authors:** Sapna Chhabra, Victoria Mochulska, Carina B. Vibe, Anubhuti Anushree, Kristina S. Stapornwongkul, Thomas Thumberger, Joachim Wittbrodt, Paul François, Alexander Aulehla

**Author notes:** These authors contributed equally, ordered alphabetically (Sapna Chhabra, Victoria Mochulska, Carina B. Vibe).

## Abstract

How organisms develop in dynamic environmental conditions is a fundamental question. We asked how day-night temperature cycles impact embryonic axis elongation and segmentation, itself a cyclic process linked to the segmentation clock, using the Japanese rice fish medaka. We developed an unbiased dimensional reduction approach, based on Singular Value Decomposition (SVD), to reliably identify the dynamic modes of segmentation clock oscillations across all temperature conditions. We reveal that the two major dynamic modes show opposite temperature sensitivities: while the temporal oscillation (mode 1) varies strongly with temperature, the spatial phase gradient (mode 2) appears largely temperature invariant. In addition, we found developmental parameters with intermediate, sub-scaled temperature responses, such as axis elongation. We used theoretical modeling to understand how dynamic modes emerge from the underlying local oscillation dynamics and axis elongation. We then exposed embryos to circadian and ultradian temperature cycles to reveal dynamic response patterns of oscillations and axis elongation, and found how these responses are integrated into morphological features. Combined, our theoretical-experimental results support a model in which the dynamic integration of temporal (i.e. segmentation clock related) and spatial (i.e. axis elongation) processes, in particular their sub-scaled temperature response patterns, quantitatively compensate each other to yield a robust, temperature-invariant axis patterning outcome.

## INTRODUCTION

One of the defining characteristics of living organisms is their ability to integrate and respond to their dynamically changing environment. The integration of environmental cues is particularly important during the earliest stages of life, i.e. during development. It is hence of fundamental importance to understand how the environment, such as changes in temperature or nutrition, influence developmental processes and ultimately, the resulting phenotype. In regard to temperature, it is clear that the rate of developmental progression is highly sensitive to changes in temperature [1]. In some context, the effect was described as being a uniformly scaled across development, potentially suggesting a common, global regulation of all processes [2]. However, at the phenotype level the effect upon changes in temperature is very much context dependent. Temperature sensitive as well as temperature compensated traits are found, even within the same organism. For instance, medaka (Oryzias latipes), a small fresh-water fish used in this study, has been reported to be mostly robust to temperature changes in terms of its overall body morphology and patterning, while at the same time, showing temperature-dependent sex determination with clear impact on male-female ratios [3–5].

To study how naturally occurring changes in temperature impact processes at molecular and signaling level, and how these responses are integrated across multiple levels or organization, yielding phenotypic consequences, we study body axis segmentation in medaka embryos.

In their natural habitat, i.e. shallow-water rice fields, Oryzias/medaka (*Oryzia* lat. rice) experience temperature changes at two timescales, seasonal and day-night temperature changes. Extensive seasonal variation ranges from 0°C to 35.9°C during the whole year, and 10.0°C to 36°C during the spawning season [6]. Remarkably, individual embryos also encounter significant periodic changes in temperature over several days during their development into hatchlings. These cycles in water temperature are the consequence of the diurnal day-night temperature fluctuations and the relatively shallow water column, measured to vary on average between 21 and 27 C over 24 hours during the month of June (see Supplement and [7]). The environmental cycles in temperature are expected to speed and slow down, respectively, developmental progression and hence body axis segmentation, though so far, to the best of our knowledge, these responses and phenotypic consequences of temperature cycles have not been studied.

Body axis segmentation, which results in the formation of somites, the precursors of the vertebral column, is in itself a periodic event, which repeats approximately every 80 minutes (at 27 degrees) in medaka embryos. Its rhythm is linked to an underlying oscillatory gene activity network, composing the vertebrate somite segmentation clock [8–10]. It involves oscillations in Notch, Wnt and Fgf signaling pathway activities, and underlies the species characteristic rhythm of somite formation, e.g. 2 hours in mouse, 30 minutes in zebrafish, 6 hours in human embryos [9, 11]. While differences in network composition have been found across vertebrates [12], interestingly, highly conserved commonalities at the level of oscillations dynamics have been identified across all vertebrate species studied. These include, on one hand, a gradual slowing down of oscillations as cells mature, which effectively establishes a period gradient that runs along the body axis [13–16]. Second, cells are coupled and synchronized to neighboring cells via direct cell-to-cell contact and Notch signaling [17–19]. Combined, these features underlie the emergence of spatiotemporal wave patterns that are seen traveling along the embryonic body axis from posterior to anterior, a characteristic found across all vertebrates studied.

Given these highly complex dynamics, which in addition are integrating additional cues, such as spatial signaling gradients and axis growth, our aim of this study was to establish a theory-experimental approach to systematically and quantitatively analyze the response patterns to changes in temperature. We hence developed an analysis strategy based on Singular Value Decomposition (SVD)[20], inspired by machine learning, in order reduce dimensionality with the aim to reveal the essential modes of oscillation dynamics. Strikingly, only two SVD modes (e.g. temporal oscillation and spatial phase gradient) are sufficient to faithfully capture the data. Importantly, by performing SVD across different temperature conditions, we revealed distinct quantitative responses of both dominant modes to changes in temperature. In addition, at the quantitative level, a third temperature response pattern was detected at the level of local oscillation dynamics and elongation dynamics.

We used theoretical modeling to understand how dynamic modes emerge from the underlying local oscillation dynamics and axis elongation and tested experimentally the dynamic response to circadian and ultradian temperature cycles. Combined, our theoretical-experimental results support a model in which the dynamic integration of several sub-scaled temperature responses quantitatively compensate each other to yield robust, temperature-invariant patterning

## RESULTS

### Developmental rate increases with temperature in medaka embryos

To examine the effect of temperature on developmental progression, we cultured embryos under constant (chronic) and cycling temperature conditions, covering the range of temperatures observed in the natural habitat of Medaka (Fig. 1A). For constant conditions, we selected the temperatures that ensured 100% hatching efficiency in the laboratory environment (17– 32 °C, Fig. 1C). For cycling conditions, we used a 24-hour temperature profile (range 21–27 °C; mean 24 °C) that reflects typical daily fluctuations observed during the breeding season ([7], Fig. 1A,B).

**Figure 1.**
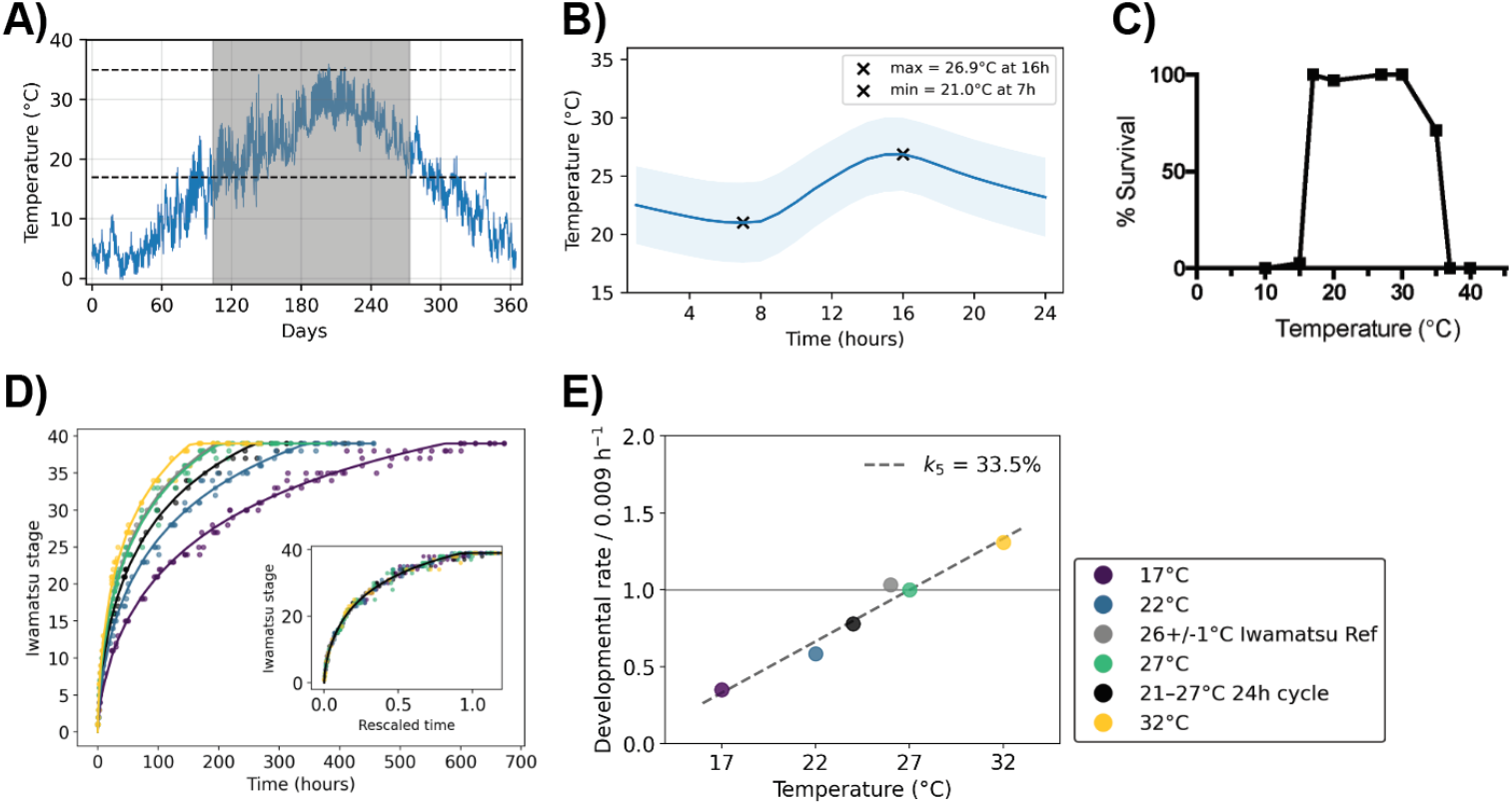
Developmental progression in different temperature conditions. **A-B)** Hourly temperature measurements taken from an outdoor medaka breeding facility in Okazaki during 2018 as a proxy to show the extent of seasonal and diurnal variation medaka might experience in the wild [7]. **A)** All data for 2018, Spawning season highlighted in gray. Dashed lines delineate the range of temperatures found to be permissive for embryo survival in the laboratory conditions (C) **B)** Average daily variation for the month of June, shown as mean± standard deviation. **C)** Percentage of embryos that successfully hatched when raised at the indicated temperature in the laboratory **D)**Developmental progression quantified by Iwamatsu staging of embryos cultured at different constant temperatures between 17-35°C (shown in colors), or during circadian variation between 21-27°C (shown in black). Inset shows the same curves in rescaled time. Experiments in C) and D) were carried out by fertilizing groups of at least 9 embryos on N separate days where N = 5, 4, 7, 11, 7 for 17°C, 21-27°C, 22°C, 27°C, 32°C respectively. **E)** Developmental rate at different temperatures. Dashed line indicates the linear fit. Legend labels for D) and E) indicated on the right side of panel E.

To quantify developmental rate, we placed embryos in different temperature conditions immediately after fertilization, and then staged them according to the Iwamatsu staging system [21] at regular intervals until they reached the hatching stage (stage 40). As expected, embryos cultured at higher temperatures developed faster and reached the hatching stage earlier than embryos cultured at lower temperatures (Fig. 1D). The embryos grown in 24-hour cycling temperature regime developed at a rate intermediate between those at 22 °C and 27 °C, in line with their average temperature of 24 °C. Despite these differences in absolute timing, the overall shape of the developmental trajectories remained similar, indicating that the overall sequence and relative timing of stages are preserved across temperatures. Consistent with this idea, all trajectories could be fitted as a function of time and collapsed onto a single curve by rescaling the time axis (Fig. 1E). From this fitting, we obtain a temperature-dependent developmental rate, *r*(*T*), which quantifies the speed (inverse timescale) of embryonic development at each temperature. Across the tested range of temperatures, this rate increases approximately linearly with temperature. We therefore fit this trend with a linear model,

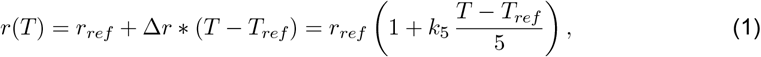

where *r*_*ref*_ = *r*(27°*C*) is the developmental rate at reference temperature *T*_*ref*_ = 27°*C*, ∆*r* is the slope of the linear fit, and 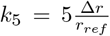 is the change in developmental rate over a 5°C interval relative to the rate at the reference temperature, 27°C. We refer to *k*_5_ as the linear temperature sensitivity coefficient. From this fit, we obtain *k*_5_ = 0.335 (Fig. 1F), indicating that a 5°C increase in temperature raises the developmental rate, in absolute terms, by 0.335 *× r*_ref_, or 33.5% of the rate at 27°C. As the model is linear, each 5°C increase produces the same absolute increase in developmental rate, whereas the fractional increase in rate varies at every temperature.

### An endogenous Her7 segmentation clock reporter for quantifying oscillations and axis elongation dynamics

To enable quantitative, time-lapse imaging measurements of segmentation clock oscillations and axis elongation, we first generated a her7–mVenus knock-in fusion reporter line. We tar-geted the endogenous Her7 locus using a CRISPR–Cas9 strategy. Validation of this reporter line showed that the Her7–mVenus protein exhibited robust oscillations in the presomitic mesoderm (PSM) (period at posterior tail bud 80 min., at 27°C,) allowing real-time quantification of the medaka segmentation clock for the first time (Fig. 2A-B and Fig. S1). Additionally, we found evidence that the reporter knock-in allele is functional. Somite development, i.e., polarity and vertebral length were similar in embryos homozygous for the Her7–mVenus allele compared to wild type embryos (Fig. S1). In contrast, when we disrupted her7 allele using CRISPR KO, we found impaired, irregular somite polarity and vertebral morphology, indicating that Her7 is a functional component of the segmentation clock network in Medaka (Fig. S1).

**Figure 2.**
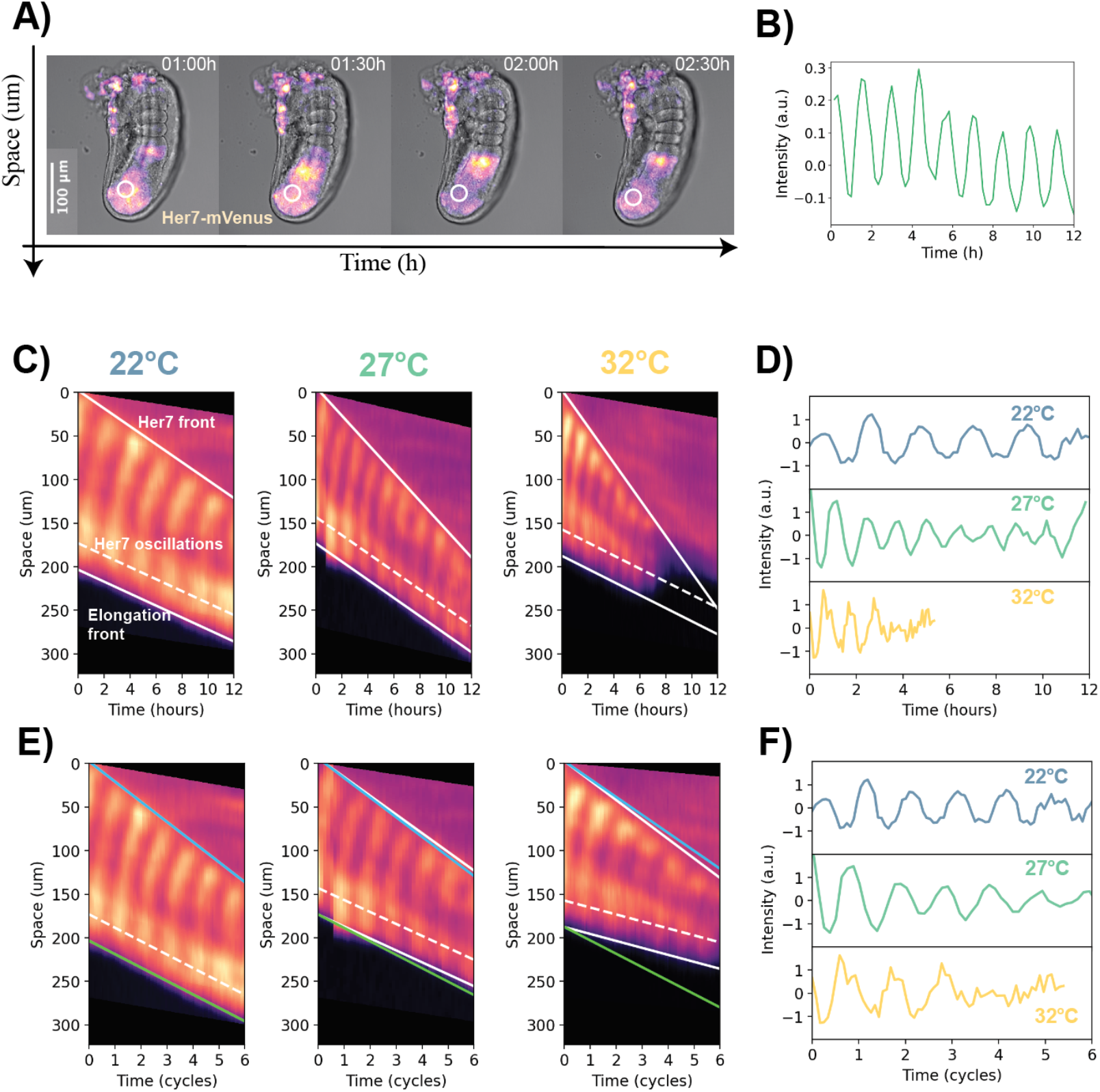
An endogenous reporter of Her7 to examine segmentation clock and axis elongation dynamics. **A)** A tail explant with Her7 reporter showing a Her7 wave propagating anterioirly from the tip of the tail. **B)** Normalized Her7 intensity profile over time in the circular ROI (region of interest) shown in A) . **C)** Representative Her7 intensity kymographs at 22C, 27C and 32C **D)** Normalized Her7 intensity profile over the dashed line in each of the kymographs in (C). **E)** Intensity kymographs in C) rescaled by the Her7 oscillation cycles. **F)** Normalized Her7 intensity profiles over the dashed lines in in the rescaled kymographs shown in (E) .

Using this validated knock-in clock reporter her7-mVenus line, we examined the temperature dependence of axis elongation and segmentation clock dynamics. To this end, we performed live imaging of her7–mVenus using embryo tail explants, at constant temperatures of 22°C, 27°C, and 32°C. We represent the dynamics of clock oscillations, traveling waves and axis elongation dynamics in the form of intensity kymographs (Fig. S3).

Qualitatively, the oscillation frequency and axis elongation increased with temperature, as expected (Fig. 2C-D). To obtain a more quantitative analysis and test whether these processes scale proportionally, we rescaled the 22°C, 27°C and 32°C kymographs, normalizing the time axis to the Her7 oscillation frequency in the posterior. Comparing the normalized data shows that axis elongation changes in a quantitatively different manner compared to segmentation clock frequency in different temperature conditions(Fig. 2E-F). These findings provided the first evidence that the temperature scaling response of different developmental processes is not uniform.

### Mode decomposition of spatio-temporal oscillation dynamics

To precisely quantify the temperature dependence of the spatio-temporal segmentation clock oscillations across multiple temperature conditions, we developed an unbiased dimensional reduction approach. To this end, we used Singular Value Decomposition (SVD) to analyze time-lapse imaging data. Our inspiration comes from theoretical machine learning, where SVD has been shown to capture semantic information about data in a hierarchical way [20]. In our present context, SVD decomposes phase kymographs based on the time-lapse measurements into a sum of dynamical modes. Each mode is itself a product of a spatial and a temporal dependency (Fig. 3A-C, Fig. A3) such that

**Figure 3.**
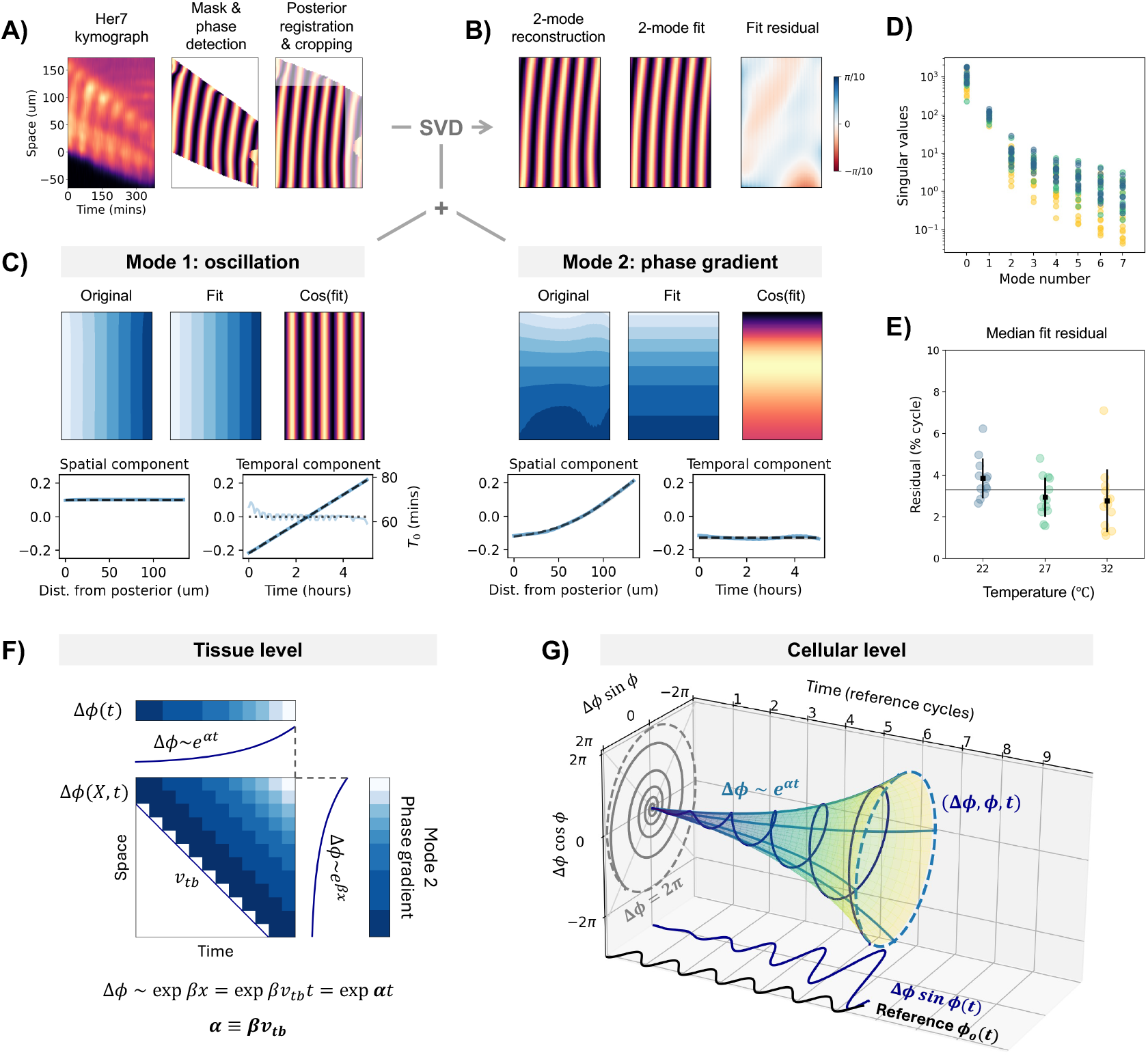
Mode decomposition. **A)** Schematic of the data preprocessing. Fluorescence images are converted into phases, then a rectangular region is cropped to perform Singular Value Decomposition. **B)** Original data are further compared to Mode 1+2 reconstruction and corresponding fit for a specific example, the right panel displays the residuals after fitting, showing a typical agreement between actual and reconstructed phase around 5% of a cycle (i.e. *π/*10) **C)** Extraction and fitting of the first two SVD modes. Mode 1 corresponds to a spatially uniform oscillation, while Mode 2 encodes a spatial phase gradient constant in time. We also compare the raw modes to linear temporal and spatial sigmoid fits for Mode 1 and Mode 2 respectively (dashed lines). **D)** Singular values spectrum of the first 8 modes in chronic temperature samples. **E)** Median (over kymograph pixels) residual of the 2-mode fit in chronic temperature samples. **F)** Reconstructing cellular dynamics from the phase gradient (Mode 2) and axis elongation (*v*_*tb*_) in the non-moving (anterior) frame. The accumulation of phase shift ∆*ϕ*(*X, t*) is characterized by the rate *α*. **G)** Cellular dynamics represented in cylindrical coordinates (∆*ϕ, ϕ, t*).

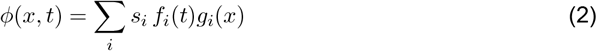

where the *f*_*i*_ and the *g*_*x*_ are the orthogonal spatial and temporal modes, and where the ordered *s*_*i*_ capture the ‘semantic’ weight of each mode.

Strikingly, only two modes are sufficient to capture 99% of the oscillation phase dynamics (see eigenvalue spectrum Fig. 3D). The first mode is uniform in space and grows linearly in time (Fig. 3C left). This linear temporal component can be fitted to compute the segmentation clock frequency, which corresponds to the Her7 oscillation frequency in the tailbud region, i.e.:

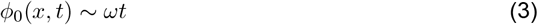

The second mode is uniform in time but increases non-linearly in space (Fig. 3C right. This spatial pattern corresponds to the spatial gradient of Her7 oscillation phase ∆*ϕ*(*x*) along the pre-somitic mesoderm, which arises due to the gradual slowing down of oscillations in cells as they shift anteriorly. The resulting nonlinear spatial phase profile can be fitted with a sigmoidal function to extract a new parameter *β*, defined as the slope of the phase gradient, such that:

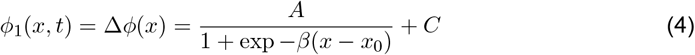

*β* quantifies how rapidly the phase changes across the tissue (see Supplement for the definition and fitting of *A, C* and *x*_0_ parameters).

### Modeling multiple time-scales of oscillation dynamics and their link to spatial phase gradients

Intuitively, we expect the spatial phase gradient to relate to the clock slowing down as cells move anteriorly. Cells are moving anteriorly with respect to the tailbud with a speed *v*_*tb*_, corresponding to the elongation speed (Fig. 3F), so that their relative position to the tail bud is *x*(*t*) = *v*_*tb*_*t* (assuming they are at the tail bud at *t* = 0). As they move anteriorly, they accumulate a phase gradient, Eq. 4 ∆*ϕ*(*x*(*t*)) = ∆*ϕ*(*v*_*tb*_*t*), which, for small enough time (or small *x* « *x*_0_) behaves exponentially :

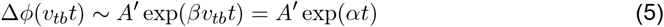

where we define *α* = *βv*_*tb*_. This parameter *α* defines a new timescale (distinct from the oscillation period), quantifying the rate of slowing down of the oscillating cells. Remarkably, these results based on SVD analysis of time-lapse quantification in medaka coincide with the prediction of a phase model, i.e. the *α*-model, previously derived from scaling experiments in mouse embryo explants [22]. Our quantifications hence reveal multiple distinct time-scales, i.e. oscillation period and oscillation slowdown, and indicate that these are general features of the segmentation clock oscillatory system.

We represent the multiple timescales during segmentation clock oscillations dynamics as ‘time cones’ (Fig. 3G). A cell oscillation is represented with a helicoidal line in 3D

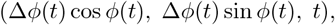

which we parametrized by the time since tailbud exit *t* and the phase of the tailbud (i.e. reference) oscillator *ϕ*_0_(*t*) = *ωt*. This allows to visualize the phase shift that a cell has accumulated over time ∆*ϕ*(*t*) as the envelope of the cone, and its current phase *ϕ*(*t*) as the position on the cone.

The accumulated phase shift ∆*ϕ*(*t*), which behaves as *e*^*αt*^, combined with the elongation dynamics (*v*_*tb*_), underlies the build-up of the spatial phase gradient that spans the PSM ∆*ϕ*(*x*) ∼ *e*^*βx*^. Because *x* = *v*_*tb*_*t*, the relation of oscillation slowing down, elongation dynamics and shape of spatial phase gradient are captured with

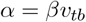

(Fig. 3F).

In summary, the quantifications combined with theoretical modeling hence allow to identify multiple timescales of oscillation dynamics within the PSM and integrate it with axis elongation to the resulting spatial phase gradient.

### Temperature sensitivity of dynamic modes and of axis elongation

After having established the SVD-based approach and with a model of how timescales, elongation dynamics and spatial phase gradient are linked at hand, we next analyzed the temperature response of the dynamics modes and body axis segmentation, Fig. 4A. Our analysis included the temperature response of following measured parameters: *ω, β, v*_*tb*_, and the speed of the Her-7 front propagation *v*_*f*_ . We combined those parameters to further compute *α* = *βv*_*tb*_, *α*_*c*_ = 2*π α*/*ω* corresponding to the slowdown per cycle, and finally, two length-scales: *λ*_*f*_ = 2*π v*_*f*_ / *ω*, the pattern wavelength at the Her-7 front (a proxy for segment size), and *λ*_*tb*_ = 2*π v*_*tb*_/ *ω*, elongation per cycle. Within this physiological temperature range, all those parameters showed an approximately linear temperature dependence (Fig. 4B). For each parameter, we then computed their respective temperature sensitivity coefficients, *k*_5_ (analogous to Eq. 1). For instance, the segmentation frequency has *k*_5_ = 36.1%, corresponding to an increase of 36.1% *ω*(27°*C*) per 5 degrees. In order to evaluate whether the temperature response is uniform across all the processes captured, we rescaled all *k*_5_s with respect to the temperature coefficient of the segmentation clock frequency, 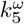 (Fig. 4C).

**Figure 4.**
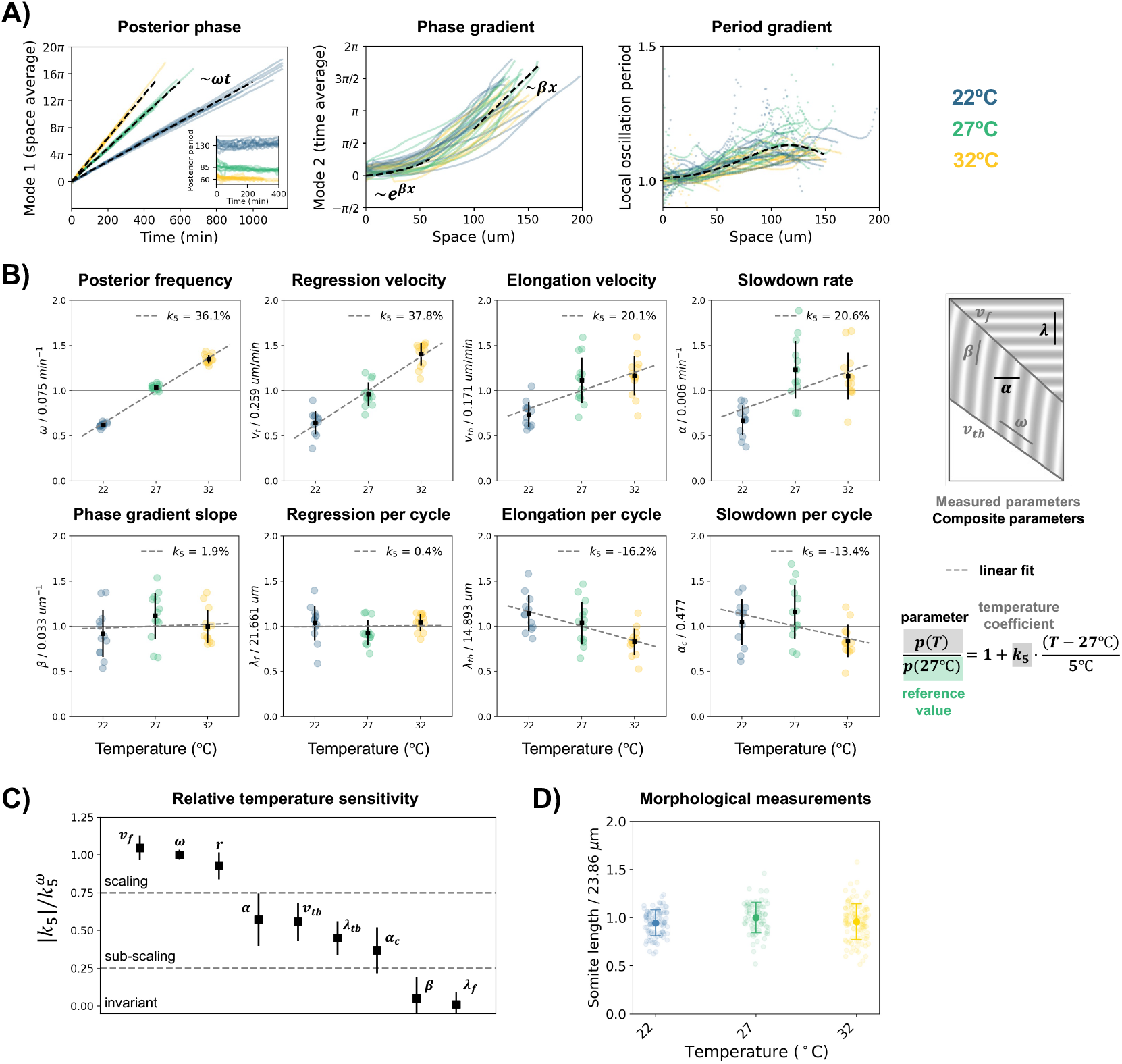
Scaling of system parameters with temperature in chronic temperature conditions. **A)**SVD modes in all chronic temperature samples, color indicates temperature. Modes were scaled by the singular value and shifted to start at 0. The slope of mode 1 defines the frequency of oscillations in the posterior. The fit of mode 2 defines the spatial gradient steepness *β*. Local oscillation period is calculated from the phase gradient and elongation velocity, shown rescaled to the posterior period. **B)** Temperature responses of system parameters. Each temperature dependence is fitted with a linear function, defining the temperature coefficient *k*_5_. **C)** Temperature coefficients *k*_5_ of different parameters relative to the temperature coefficient 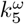 of the clock frequency *ω*. Three groups of parameters are defined based on this relative sensitivity: scaling, sub-scaling, and invariant. **D)** Morphological measurements of nascent somite length at different temperatures, pooled from different samples and somite stages.

Interestingly, this analysis revealed distinct temperature responses for different processes and parameters (Fig. 4C). We defined three categories: (a) processes that showed the same quantitative response to temperature as the clock (i.e. ‘scaling’ parameters such as the global developmental rate (*r*), as well as the speed of the regressing Her-7 front *v*_*f*_); (b) a group of processes showing a quantitatively distinct response (i.e. ‘sub-scaling’), i.e. axis elongation and clock slowing down; and finally (c) processes and parameters that we found to remain invariant across different temperatures (‘invariant’, i.e. phase gradient and length of segments). Notably, we found that axis elongation changes relatively less compared to oscillation frequency when temperature is altered, i.e. the scaling sensitivity factor k for *v*_*tb*_ shows lower values (Fig. 4C, i.e. ‘sub-scaling’). This quantification is consistent with the visual observations in Fig. 2E, where rescaled elongation speed appears much smaller at 32C compared to 22C . Such difference in temperature scaling between the clock period and axis elongation suggests that these processes are independently controlled, reflecting their modular nature [23].

At the same time, we found that several parameters are temperature invariant *k* ∼ 0, which includes the phase gradient shape, quantified by *β*. Our model allows to interpret temperature invariant *β* as resulting from the ‘sub-scaling’ response of oscillation slowing down and axis elongation speed (i.e. *α* = *βv*_*tb*_). Their combined response to temperature, which suggests the existence of an active process connecting the clock slowing-down (*α*) to the elongation speed (*v*_*tb*_), effectively results in the invariant phase gradient shape *β* that we found across all temperature conditions. In addition to the temperature-invariant phase gradient, this category of parameters also included the segment pattern wavelength at the Her-7 front, *λ* = 2*π v*_*f*_ / *ω*, which is a proxy for somite size.

In agreement with this result, we found that direct morphological measurements of nascent somite size showed that the average somite size indeed remained invariant across temperatures, Fig. 4D.

Combined, our analysis thus revealed quantitatively distinct temperature responses of dynamical parameters, e.g. clock frequency vs elongation speed and clock slowing down. These distinct responses combine and are integrated to yield overall invariant dynamical features and morphological outcome, i.e. an invariant emerging spatial phase gradient and robust average somite sizes across temperatures, respectively.

### Temperature cycles reveal dynamic integration of modular responses of segmentation clock and axis elongation

To determine if the temperature response of the segmentation clock and axis elongation can be further uncoupled, we subjected embryos to periodic temperature changes, including the natural 24-hour temperature cycles experienced by Medaka embryos in the wild and higher frequency harmonics (12- and 6-hour cycles). We then quantified the dynamic response of the segmentation clock frequency and elongation velocity (Fig. 5A-B).

**Figure 5.**
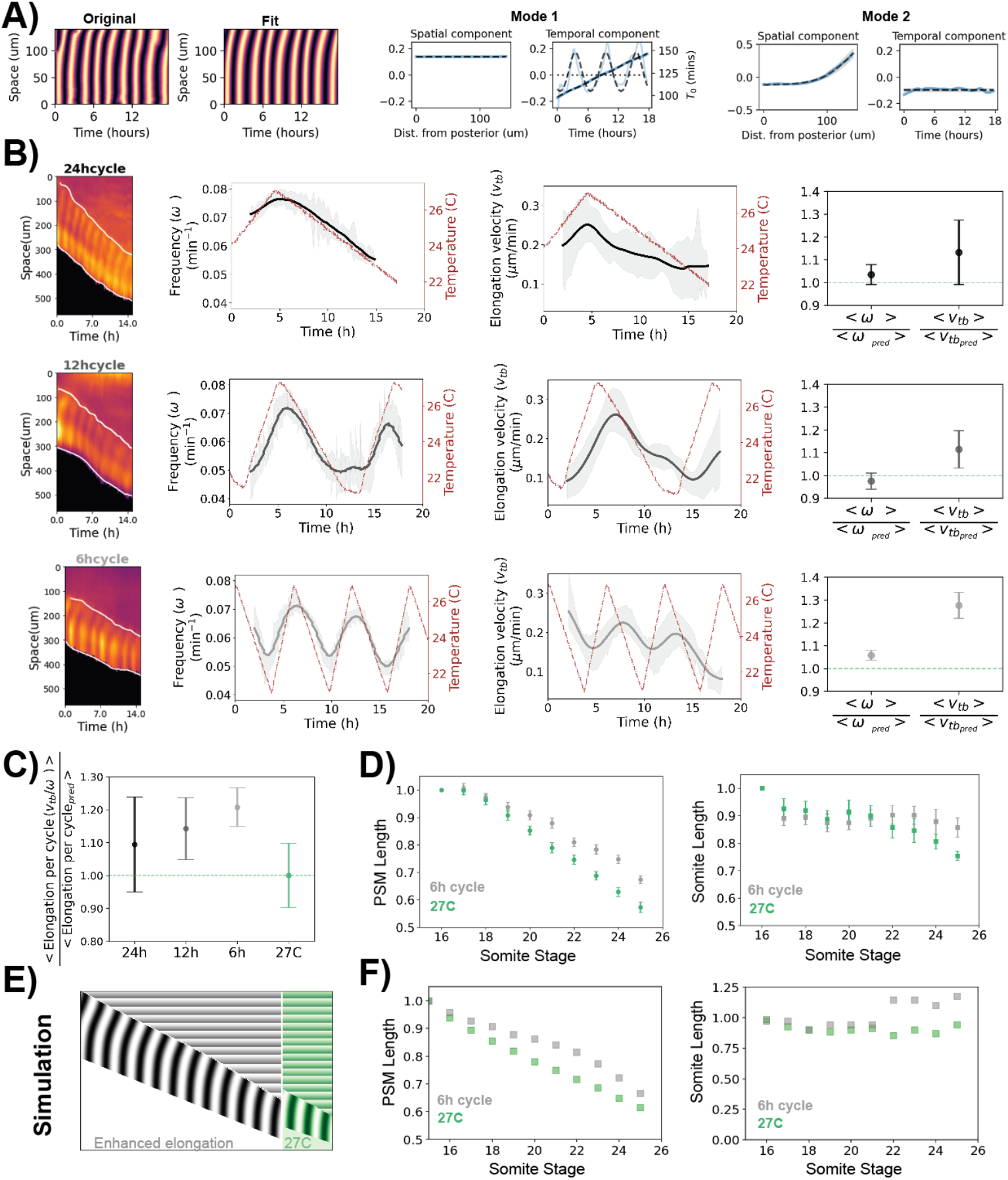
Dynamics of segmentation clock and axis elongation in cycling temperatures. **A)** SVD analyses of an intensity kymograph in 6h temperature cycles. Temporal component of Mode1: clock oscillation, spatial component of Mode2:phase gradient. **B)** Dynamics of clock frequency and elongation velocity in different cycling temperatures. *Left:* representative kymographs, *Middle:* Temporal profiles of segmentation clock frequency and elongation velocity. Solid lines represent mean value across all samples. Dashed region represents standard deviation across samples. *Right:* Experimental average relative to predicted average values obtained from data in chronic experiments. Predicted values obtained according to the equation, *p*(*T*) = *p*(27*C*) * (1 + *k* * (*T −* 27*C*), where *p*(27*C*) is the value of the relevant variable at 27C and k represents its thermal sensitivity. *r*(27*C*) = 0.078, 0.191 and *k* = 0.08, 0.068 for segmentation clock frequency (*ω*) and elongation velocity (*v*_*tb*_) respectively. For all datasets, *N* > = 2 and *n* > = 9. **C)** Axis elongation per segmentation clock cycle in different cycling conditions (24h, 12h, 6h cycles) and 27C chronic condition. **D)** PSM and somite length progression in 6h cycling and 27C chronic temperatures. These measurements are obtained from tail morphology. Refer Fig S7. For both datasets, *N* > = 2 and *n* > = 9. **E)** Kymographs simulated with alpha model, comparing chronic temperature condition (green) with a condition where elongation is enhanced similar to that observed in different cycling conditions (gray). **F)** Simulated PSM and Somite length progression.

In all conditions, we found that the segmentation clock frequency quickly adapts to temperature, closely following the temperature cycles: it increases and decreases almost in phase with temperature, even for 6-hour temperature cycles. We then compared the average frequency observed in each cycling condition with the frequency predicted from the chronic experiments at the mean temperature of the cycle. Remarkably, we found that the segmentation clock frequency oscillates around its predicted value. This suggests that the segmentation clock precisely mirrors the temperature cycle, adjusting its period to the one predicted from chronic conditions, with a time scale much faster than our highest cycle frequency (6 hours) (Fig. 5B).

In contrast, elongation velocity during temperature cycles exhibits a markedly different behavior. Although elongation velocity also varies with temperature, we observed a clear delay of 2 hours upon changes in temperature. Furthermore, the elongation speed averaged over these cycles is about 20% higher than the elongation speed predicted from the corresponding temperature in chronic conditions. This indicated that cyclic changes in temperature enhance elongation relative to constant-temperature conditions (Fig. 5B). Mechanistically, we found evidence for an entrainment effect of cyclic temperature changes on elongation dynamics. First, we found that elongation exhibits periodic changes in its dynamics, with a period of 5 h across conditions (Fig. S5A-C). Second, in the presence of temperature cycles, we see that elongation cycles become more synchronous across different embryos, a hallmark of being entrained by an external rhythm (or zeitgeber) (Fig. S5D).

As a consequence of the differential temperature response of clock and elongation velocity, there is a 15% increase of elongation per clock cycle when temperature conditions are cyclic (Fig. 5C). To determine how the increase in elongation per segmentation clock cycle impacts morphological outcome, we measured somite and PSM length upon exposure of embryos to 6h cycling temperature conditions. We found that compared to measurements done in embryos cultured at constant temperature, 6-hour temperature cycles indeed led to an increased PSM length, matching the increased elongation velocity. Interestingly, somite length first remained unaltered compared to control embryos, but with a delay of ∼ 3 cycles, we noticed that somite sizes increased, in parallel to an increased PSM length. These results imply that the immediate response to 6-hour temperature cycles is an increase in elongation velocity per clock cycle, which leads to a delayed change in somite size over time (Fig. 5D).

To understand the underlying mechanism, we turned to mathematical modeling (Fig. 5E, Fig. A6). In our theoretical framework, given an invariant beta, a change in elongation velocity *v*_*tb*_ is directly linked to a change in the slowdown of oscillations (*α*), consistent with the relation *α* = *βv*_*tb*_. In the model, we assumed that the change in *α* upon changes in *v*_*tb*_ occurs only *locally* in the posterior PSM, the region driving elongation. The resulting model simulations revealed that with an increase in elongation *v*_*tb*_ and a local feedback on *α*, indeed the PSM length increases immediately, while somite size initially remains unchanged, matching experimental findings. Only with a delay, i.e. after the time required for posterior cells in which *α* was altered to advect into the anterior region, the simulations show that somite size increases again closely matching experimental findings (Fig. 5F).

The theoretical framework hence provides a concrete possible mechanistic explanation and insight into the dynamic feedback regulation between elongation, the multiple time scales of oscillation (i.e. the period and independently, the rate of slowing down *α*) and segment size.

More generally, these results demonstrate that the segmentation clock and axis elongation form two distinct and independently regulated modules, each with a distinct response pattern, in regard to timing, magnitude and amplitude of response to temperature (Fig. 5, Fig. S6). Together with theoretical modeling, we hence reveal a possible mechanism and dynamic feed-back underlying the ability of medaka embryos to show robust, proportional axis segmentation under environmental changes in temperature.

## DISCUSSION

We have established Medaka as a system to understand how changes in temperature, in particular naturally occurring diurnal temperature cycles, are integrated into the cyclic developmental process of body axis segmentation. We show that temperature does not act through a single global scaling of segmentation. Instead, the segmentation clock and axis elongation respond as distinct modules with different thermal sensitivities, whose dynamic integration preserves temperature-invariant segment patterning. This reveals how developmental plasticity in clock and morphogenetic dynamics can be coupled to phenotypic robustness.

This insight was made possible by combining theoretical, analytical and methodological advances. First, we developed a novel endogenous knock-in her-7 clock reporter medaka line to quantify spatiotemporal clock and axis elongation dynamics under various chronic and cyclic temperature conditions, in real-time.

Critically, we combined these measurements with Singular Value Decomposition (SVD), a dimensional reduction method related to deep learning [20], which enabled a reliable, quantitative analysis of dynamical modes within the time-series data.

Two dominant, interpretable modes were sufficient to accurately describe the segmentation clock and wave dynamics, in space and time. The first mode is to a global (i.e. position-independent) phase increase corresponding to the clock oscillation. This enabled us to rigorously define the segmentation clock period across temperature conditions, including during temperature cycles. The second mode characterizes a time-invariant spatial phase gradient along the body axis, which is characterized by a slope parameter (*β*). Combining the SVD analysis and theoretical modeling also allowed us to precisely identify how the two dominant modes are linked to local oscillations slowing down *α* and axis elongation *v*_*tb*_.

Remarkably, we found that dynamic modes, local oscillation slowing down and axis elonga-tion all have distinct temperature sensitivities: while the segmentation clock frequency (mode 1) varies strongly with temperature, we found that the phase gradient (mode 2), appears largely temperature invariant.

The invariance of the spatial phase gradient might appear counterintuitive, because this gradient is generated by processes that are themselves temperature-sensitive. Oscillations slow down in PSM cells as the axis elongates - both processes change with temperature, but neither followed the scaling of the global clock frequency or overall developmental rate. Instead, both displayed weaker, sub-scaled responses. The phase gradient invariance can be explained based on these two sub-scaled processes compensating each other quantitatively across all conditions. In the model, the invariant phase gradient (*β* parameter) is formalized by the relation

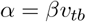

This mathematical constraint raises the biological question of whether elongation and clock slowing are functionally coupled, rather than merely correlated. Directly testing such coupling will require the ability to experimentally tune elongation speed or clock slowing, which remains to be achieved in future studies. However, our experiments under cycling temperature conditions provide indirect evidence for functional coupling between these processes.

Cycling temperatures enhanced axis elongation relative to elongation at the same mean temperature. The enhanced elongation observed in cycling temperatures transiently affected PSM length and was dynamically integrated into delayed changes in somite length. These morphological changes are captured by a dynamic feedback regulation model in which changes in elongation dynamics not only affect cell advection directly, but also lead to changes in clock slowing dynamics (Figure 5D–F; Figure A6).

Conceptually, temperature invariance therefore originates from the integration of temporal *and* spatial temperature effects across distinct developmental processes. This is in stark contrast to another well-studied biological clock, i.e. the circadian rhythm, in which compensation is achieved by various mechanisms (depending on the organisms) *within* the clock itself [24], enabling circadian rhythms to function with a temperature-invariant frequency[24]. In contrast, during axis elongation, the embryonic rhythm itself is plastic, and only in combination with axis growth, the function of this integrated embryonic rhythmic system remains temperature-invariant *in space*. To our knowledge, a coupled temporal–spatial temperature compensation mechanism that operates across different processes has not been previously described. We term it *chronomorphic compensation*.

Overall, our work reveals that temperature regulation of development is more modular than previously appreciated, with different developmental processes showing distinct temperature sensitivities. While some processes, such as segmentation clock frequency, follow the global scaling of developmental progression, other processes, such as axis elongation, scale quantitatively differently. In this context, zebrafish provide an informative comparison, as previous work suggested that segmentation clock dynamics and axis elongation scale uniformly with temperature in this species [25]. Thus, the extent to which developmental processes are thermally coupled may itself vary across species, providing a potential mechanism for the evolution of species-specific developmental responses to temperature.

In addition and unexpectedly, the cyclic temperature experiments also revealed that temper-ature cycles enhance axis elongation, an effect that is temperature independent but attributed to the cyclic nature of changes. Hence, elongation is enhanced when temperature changes periodically, compared to elongation at the equivalent average temperature. This effect was seen at all cyclic temperature regimes and was most pronounced in 6-hour temperature cycles. The underlying mechanisms remain to be revealed. These results point towards the possibility of resonance effects between environmental and developmental rhythms, a fundamental question that requires future investigations.

Our findings add a new perspective to a century-old discussion on how the temperature sensitivities of developmental rate and growth differ [26, 27]. In temperature-size rule models, the tendency of organisms to mature at smaller sizes at higher temperature is explained by the developmental rate showing greater temperature sensitivity than growth [28, 29]. Consistent with this, we found that developmental progression and segmentation clock frequency have higher temperature sensitivity than axis elongation (Figures 1E and 4C), suggesting that differential thermal sensitivity between developmental timing and tissue growth is already apparent during embryogenesis.

However, our data based on more granular quantitative analysis of developmental processes and their underlying modules revealed that also *within* development, different sensitivities are visible (Figure 4C). In other words, differences in temperature sensitivities are not only seen between developmental rates and growth, but rather, these differences also occur within developmental processes. Accordingly, our quantifications revealed multiple-time scales within the segmentation clock, i.e, the clock period and the rate of oscillation slowdown (Figure 3G), each showing a distinct temperature sensitivity.

These distinct temperature response patterns emphasize the modular nature of development and provide a template on which evolution of feedback regulation can act to balance developmental plasticity with phenotypic robustness.

## METHODS

### Medaka Maintenance

Medaka were kept in a recirculating system at 27°C, with a 14:10 light/dark cycle. Wildtype strains were a kind gift from the Wittbrodt lab, derived from the Cab inbred strain, in crossed to at least the 64th generation. Fish were housed in the animal facility at EMBL following the guidelines of the European Commission, revised directive 2010/63/EU and AVMA guidelines 2007 and under veterinarian supervision. Approval for animal experiments was obtained from the EMBL Institutional Animal Care and Use Committee (project codes: 21–001_HD_AA, 25-014_HD_AA).

### Generation of Her7 mVenus and Her7 knockout lines

Endogenously tagged medaka Her7 fusion protein line and the Her7 knockout line were generated using CRISPR-Cas9. sgRNA candidates were designed using the online sgRNA target predictor tool CCTop [30]. Targets were verified by sequencing of the targeted region in our wild-type medaka colony, and synthesized using the Megshortscript kit (ThermoFisher, AM1354) followed by RNA cleanup using the RnEasy minElute cleanup kit (Qiagen, 74204).

For cloning of donor plasmid for the Her7-mVenus line, homology arms (647 and 641 bp)) were chosen to match the sequences flanking either side of the stop codon. To protect donor integrity and prevent re-cutting after homologous repair, sgRNA sites were mutated in the donor construct by mutagenesis PCR. Donor construct was assembled using Golden GATE-way cloning [31]. Cas9 was supplied in the form of Mammalian codon optimized Cas9 mRNA (Addgene plasmid 43861) or as a Cas9 variant gifted from the Wittbrodt lab synthesized with the mMessage mMachine SP6 kit (ThermoFisher Scientific, AM1340).

The Her7-mVenus line was first created by co-injecting Cas9 mRNA, the Her7-mVenus donor plasmid and two sgRNAs (GCCAGACTCTGTGGAGGCCCTGG and TTGGTGACCGGGTCAGGGCCAGG). This resulted in a 5’ in frame insertion of the mVenus coding sequence in the desired location, but also included part of the flanking 3’ donor backbone and a region of non-homologous end-joining before the endogenous UTR, extending the locus by 3.5kb. To remove the extra insertion, a second round of CRISPR/Cas9 was performed with sgRNA TAGCAGTTATTCCATCGGCACGG, resulting in a clean, in-frame integration of mVenus. The Her7 knockout line was created by co-injecting Cas9 mRNA with two sgRNAs (CATCCCGGAAGGACTGCTTCTGG and AAAGTTACTAAAATCTCAGGTGG) targeting either side of the bHLH domain, which mediates DNA binding and dimerization [32].

Medaka embryos were injected at the one-cell stage with 150 ng/μl Cas9 mRNA, 15 ng/μl sgRNA and 5 ng/μl donor plasmid (if present) and screened by PCR for in-frame integrations. To minimize the possibility of off-target effects, both lines were also crossed out to wildtype.

### Morphological characterization of embryos and hatchlings in different temperature conditions

Embryos were collected from pre-separated breeding pairs after 15 minutes of breeding, and placed in Embryo Culture Medium [33] at different temperature conditions within 15 minutes of collection. For constant temperature conditions embryos were cultured in a standard cooling incubator (Neolab 7-1104). To achieve continuously changing temperature for the diurnal variation condition embryos were cultured using a programmable cooling incubator (Binder KT53).

In all conditions, the medium was changed every other day at a minimum. Embryos were staged according to Iwamatsu [21]. Skeletal staining was performed according to [34]. Colorimetric in situ hybridisation was performed according to the protocol in [35]. Myf5 probes were a gift from Winkler lab [36].

### Live imaging of Medaka tail explants

Embryos were obtained from crosses between wild-type Cab females and Her7-mVenus homozygous males and raised to the desired developmental stage under the respective temperature conditions. For 24 h temperature cycles, embryos were maintained in cycling conditions immediately after fertilization. In contrast, for 12 h and 6 h temperature cycles, embryos were initially raised at 27°C and only subjected to the respective temperature cycles during the imaging period. Embryos were subsequently dechorionated by first weakening the chorion with fine sandpaper, followed either by enzymatic treatment with hatching enzyme or by manual removal using forceps. After dechorionation, tail explants were generated by dissecting 2–5 somites posterior to the unsegmented presomitic mesoderm (PSM).

Dissection and imaging of tail explants was performed under two distinct experimental conditions that differed in medium composition, developmental stage, fish generation, and temperature regime. To account for these differences, experiments were assigned to one of two batches. **Batch 1** comprised experiments performed between 2015 and 2021 under chronic temperature conditions or 24 h and 12 h temperature cycles. These experiments used embryos at the 13–15 somite stage and were dissected and imaged in a CO_2_-independent base medium containing phenol red (Gibco, 18045054). **Batch 2** comprised experiments performed between 2022 and 2024 under 6 h temperature cycles and the corresponding 27°C control condition. These experiments used embryos at the 16–17 somite stage and were imaged in CO_2_-dependent DMEM/F-12 medium lacking glucose, pyruvate, glutamine, and phenol red (Cell Culture Technologies), supplemented under 5% CO_2_. For dissection, the DMEM/F12 media was supplemented with 10mM HEPES (Gibco, 15360-106) instead of CO_2_. The media composition used in Batch 2 improved long-term tail explant survival and was therefore essential for observing the effects of multiple temperature cycles in the 6 h cycling experiments. For both batches, the base medium was supplemented with 2 mM glucose (45% solution in water, Sigma, G8769), 2 mM L-glutamine (Gibco, 25030), 1% penicillin-streptomycin (Sigma, P4333), and 0.1% w/v bovine serum albumin (BSA) (Millipore, ES009-B).

Live imaging was performed in the respective media with multiple samples mounted individually either in wells of a 96-well plate (ibidi, 89606) for Batch 1 experiments or in wells of a micro-insert chamber (ibidi, 80406) for Batch 2 experiments. Imaging was carried out on a Zeiss LSM780 laser-scanning inverted confocal microscope equipped with a 20× Plan-Apochromat objective (numerical aperture 0.8). For Her7-mVenus imaging, mVenus was excited at 514 nm, and images were acquired as z-stacks of 5-10 planes, spaced 7 μm apart, at a resolution of 512 × 512 pixels and a spatial sampling of 1.38 μm per pixel. In all experiments except 32C chronic conditions, images were acquired at 10 min intervals.

Temperature was maintained and controlled on the microscope using an in-house-built incubator developed by the EMBL mechanical/electronic workshop .Temperature was monitored either in a well adjacent to the sample wells, for Batch 1 experiments or in the dish outside the microchambers for Batch 2 experiments, using a digital thermometer submerged in distilled water. For experiments involving continuous temperature changes, autofocus was controlled using custom software kindly written by the Advanced Light Microscopic Facility at EMBL Heidelberg.

Both the segmentation clock frequency and the tailbud elongation velocity were lower in Batch 2 than in Batch 1 experiments (Figure S4). To account for this difference, we calculated a correction factor that equalizes the mean values at 27 °C between the two batches. This factor was then applied to the 6-hour cycling experiments in Batch 2 before comparing them with other temperature conditions in Figure 5..

### Quantifying Her7 intensity using kymographs

Her7 intensity images were processed to remove noise by blurring the max z projected image with a gaussian filter (sigma = 4). Subsequently two alternate methods were used to register images in one sample along time and generate kymographs. The first method consisted of the *(I) Center of mass registration and kymograph generation in FIJI*. Here, sample registration of the max-Z projected fluorescence channel or the one z-slice brightfield channel was carried out with the multistackreg plugin (Brad Busse, v1.45, [37]), using the Rigid body transformation algorithm. The transformation file was saved and then applied to the other channel. The registered processed intensity image was used to create a kymograph by means of the FIJI “KymoResliceWide” function, drawing a segmented line (20 pixels wide) along the entire PSM in the direction of wave propagation.

FIJI’s Multistackreg plugin was ineffective in accurately aligning many tail explants, especially the ones where either the tail moves significantly in x-y position or the direction of tail elongation changes over the course of imaging. This effect was particularly pronounced under cycling temperature conditions. To analyze such samples, a new, anterior-registration method was developed in-house. We later discovered that this method provides a better estimate of velocities than FIJI’s center-of mass registration (Figure A4), and a correction factor was subsequently applied to the velocities obtained using the center of mass registration.

*II) Anterior Registration and Kymograph generation*: This method begins with segmentation of max-z projected Her7 intensity images using the Pixel classifier option in Ilastik [38] to obtain an embryo mask for each timepoint. Since all subsequent steps depend on the mask, it is essential to get a good mask. This step is the most time-consuming of all steps since batch processing does not work well in Ilastik, and so each sample, sometimes each timepoint, had to be segmented individually. Following segmentation, the embryo masks were skeletonized to obtain the core skeleton using the bwskel function in Matlab. This skeleton was subsequently linearly extended in both directions to get the entire midline. This step was performed using a custom-written code in Matlab. The midlines were then dilated to create a region of interest surrounding the anterior-posterior axis of the tail explant. For each position along the axis, pixel intensities within the dilated region were averaged across the perpendicular section. This yielded a one-dimensional intensity profile per timepoint, capturing the spatial distribution of Her7 signal along the anterior-posterior axis of the tail explant. This step was performed using a custom-written code in FIJI. The one-dimensional intensity profiles obtained for each timepoint were averaged within spatial bins along the anterior–posterior axis and assembled into a two-dimensional array by stacking profiles sequentially in time as columns. In this representation, the x-axis corresponds to time and the y-axis to position along the anterior–posterior axis, with profiles aligned at the anterior such that the first row represents the anterior-most position or the cut site of the tail explant. The resulting matrix was visualized as an image to generate a kymograph. Kymographs were subsequently smoothed using a mild filter to reduce noise while preserving spatiotemporal features.

### Computing Her7 oscillation phases using Wavelet and Hilbert Analyses

For all batch 1 experiments, wavelet analyses (TFApy), developed by Gregor Mönke ([39]) was used to convert intensity profiles to phase and frequency information. This tool first detrends the data to remove low frequency trends from the raw signal. The detrended signal is then subjected to a time-resolved frequency analysis by cross-correlating the signal at every time-point to a set of wavelets, which are functions with a defined frequency. This generates a spectrum where for each timepoint, the wavelets which correlate well with the signal are given a high “power” score. The power is defined as how much more likely this signal is to achieve a high correlation relative to white noise. The maximum power, periods to scan for and the cut off period for the detrending is specified by the user. To extract the period values, the maximum power ridge in the spectrum is detected.

For 6h experiments, wavelet was ineffective in detecting fast changes in frequency. Hence for Batch 2 experiments – 6h cycle and the corresponding 27C experiments, phase information was obtained by performing Hilbert analyses on the detrended signal of the intensity kymograph.

## Supporting information

ManuscriptSupplement

## Acknowledgements

We warmly thank all members of the Aulehla and François labs for scientific discussion and emotional support. We thank the Wittbrodt lab for providing us with crucial support with CRISPR methods and our wildtype medaka stock. We thank Christian Kieser from the EMBL mechanical workshop for creating and optimizing the software and hardware necessary for replicating diurnal temperature variation using the microscope incubator. We thank Aliaksandr Halavatyi from the Advanced Light Microscopy Facility at EMBL for coding the custom-made autofocus software that was used during Batch 2 experiments. We thank Christian Tischer and Arif Khan from the EMBL Imaging Center for brainstorming and helping with the anterior registration and kymograph generation analyses method. We thank Dr. Ai Shinomiya, (NIBB, Okazaki) and Professor Takashi Yoshimura (Nagoya University) for kindly sharing unpublished results and for providing the raw data used for figure 1A. We thank Prof. Kiyoshi Naruse (NIBB, Okazaki, Japan) for discussions on the project. SC was supported by an EMBL Interdisciplinary Postdoctoral Fellowship (EIPOD4) program under Marie Sklodowska-Curie Actions Cofund (grant agreement no. 847543). VM was supported by an NSERC-Create training grant in Complex Dynamics and a doctoral research grant from the Fonds de recherche du Québec #328292.This work was supported by the EMBL and the Courtois Institute, and received funding from the European Research Council under an ERC consolidator grant agreement n.866537 to AA, a New Frontiers in Research Fund Exploration Award to PF and AA, and a Natural Sciences and Engineering Research Council Discovery Award to PF.

