## Supplementary material for "Cycles upon cycles - Temperature Scaling of Medaka Development": ManuscriptSupplement

### Supplemental Text

#### Fitting developmental rate and temperature dependence of parameters

To quantify the developmental rate in whole embryos, we fit the measured Iwamatsu stage $s$ as a function of time with an activation function:

$$s(t) = \frac{At^n}{t^n + t_{1/2}^n}. \tag{6}$$

For a one-parameter fit, we fix $A = 65$ and $n = 0.65$. The developmental rate is defined as the inverse half-time: $r = \frac{1}{t_{1/2}}$. This inverse timescale is then found to depend linearly on temperature.

Kymograph analysis parameters are defined in the main text (Fig. 4) and Suppl. Text section Mode fitting. For the resulting set of parameters, we analyze the dependence on temperature using a linear fit around a reference temperature $T_{ref}$. For a parameter $p$:

$$p(T) = p(T_{ref}) + \Delta p * (T - T_{ref}) . \tag{7}$$

Equivalently, dividing by the reference value gives:

$$p(T) = p(T_{ref}) * (1 + k\ (T - T_{ref})) . \tag{8}$$

Thus, the temperature coefficient $k$ is the *normalized linear slope*, i.e. the fractional change in $p$ per degree relative to $p(T_{ref})$. For the reference temperature $T_{ref}$, we choose the middle of the experimental interval: $T_{ref} = 27°C$.

For convenience, we report the relative change per a 5-degree interval, $k_5 = 5k$. With this parameterization, we have $r(32°C) = (1 + k_5)r_{ref}$, and $r(22°C) = (1 - k_5)r_{ref}$. For example, $k_5 = 0.1$ means that $p$ increases by 10% of $p(27°C)$ when temperature increases from $27°C$ to $32°C$.

When two parameters are multiplied to produce a composite parameter, $p_{prod}(T) = p_1(T) * p_2(T)$, the resulting temperature coefficient is approximately the sum of two coefficients:

$$p_{prod}(T) = p_1(T_{ref})p_2(T_{ref}) \left[1 + (k^{p_1} + k^{p_2})\ (T - T_{ref})\right] + O\left((T - T_{ref})^2\right) \tag{9}$$

Analogously, the temperature coefficient of a ratio of parameters is approximately the difference between their temperature coefficients. Examples of such combinations of temperature sensitivities are the slowdown rate $\alpha$ (product of parameters) and the elongation/regression per cycle (ratio).

For rate parameters (developmental rate $r$, frequency $\omega$, slowdown rate $\alpha$, and velocities $v_f$ and $v_{tb}$), we also fit the temperature dependence with an allometric scaling equation and the Arrhenius equation [26]. The allometric fit is given by :

$$p(T) = aT^b = p(T_{ref}) \left(\frac{T}{T_{ref}}\right)^b \tag{10}$$

The Arrhenius equation is

$$p(T) = a \exp\left(-\frac{E_a}{k_B T}\right) = p(T_{ref}) \exp\left(-E_a \left(\frac{1}{k_B T} - \frac{1}{k_B T_{ref}}\right)\right), \tag{11}$$

with $k_B$ the Boltzmann constant, and temperature in Kelvins.

We note that for a general temperature dependence $p(T)$, the linear approximation around $T_{ref}$ can be written as :

$$p(T) \approx p(T_{ref}) \left(1 + \frac{p'(T_{ref})}{p(T_{ref})}(T - T_{ref})\right) \tag{12}$$

Thus, the linear temperature coefficient $k$ can be related to the nonlinear fits through the relative slope at $T_{ref}$, $k \approx p'(T_{ref})/p(T_{ref})$. In particular, for the allometric fit $k \approx \frac{b}{T_{ref}}$, and for the Arrhenius fit $k \approx \frac{E_a}{(T_{ref})^2}$.

The nonlinear fitting results are provided in Fig. A1. Consistent with our linear fitting, the scaling exponent $b$ and the activation energy $E_a$ are similar for parameters $r$, $\omega$, and $f$ (scaling parameters), while for $v_{tb}$ and $\alpha$ (sub-scaling parameters), $b$ and $E_a$ are around twice as low.

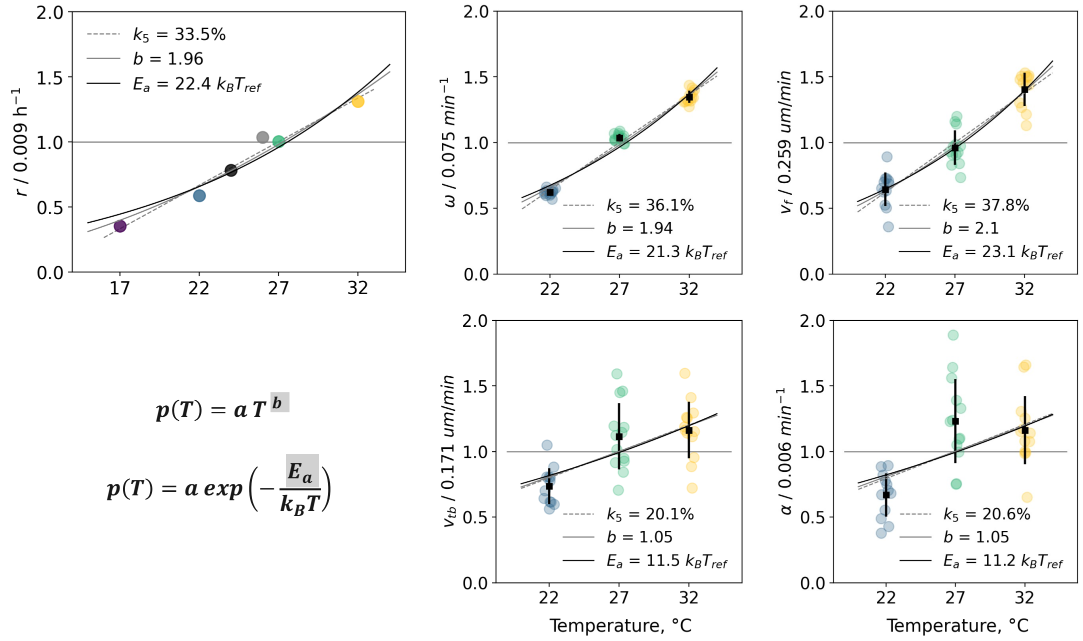


**Figure A1. Temperature dependencies of rate parameters**. Data were fitted with a linear fit (temperature coefficient $k_5$) as well as allometric scaling (exponent $b$) and Arrhenius equation (activation energy $E_a$).

#### Frames of reference

Throughout our analysis, we consider kymographs in several frames of reference. For SVD analysis and phase gradient quantification, we use the *posterior registration* frame, with the tailbud position fixed and the distance from the tailbud denoted as $x$. The space axis is directed from the posterior to the anterior.

For velocity measurements, we consider the *anterior registration* frame as the lab frame. In the anterior registration, the position of the cut (anterior edge of the tail explant) is fixed, the tail elongates with velocity $v_{tb}$, and the Her7 front moves through the tissue with velocity $v_f$. We denote the space coordinate in this frame as $X$. The tailbud coordinate is then $X_{tb}(t) \sim v_{tb}t$.

Although velocities $v_{tb}$ and $v_f$ are defined in the lab (anterior) frame, they can be conveniently measured in the *center of mass* (CM) frame. For chronic temperature kymographs, we directly measured velocities in the CM frame and calculated the anterior frame velocity by adding the CM movement with respect to the anterior edge:

$$v_{tb} = v_{tb}^{CM} + v_{CM}; \quad (13)$$

$$v_f = v_f^{CM} + v_{CM}. \quad (14)$$

To calculate the CM velocity in the anterior frame $v_{CM}$, we considered the tail explant as an elongating trapezoid, with the anterior and posterior widths approximately fixed and only the length (height of the trapezoid) increasing linearly in time with speed $v_{tb}$. With those assumptions, the center of mass of the trapezoid moves proportionally to the elongation: $v_{CM} = cv_{tb}$, where the coefficient $1/3 \leq c \leq 1/2$.

We measured $c$ experimentally by registering $N = 10$ chronic 22° C and 27° C samples in the anterior frame, directly measuring $v_{tb}$ and comparing to $v_{tb}^{CM}$ (Fig. A2 A). We found that $v_{tb} = v_{tb}^{CM} + v_{CM} = \frac{1}{1-c} v_{tb}^{CM} = 1.5 v_{tb}^{CM}$, or

$$v_{CM} = 0.5 v_{tb}^{CM} \quad (15)$$

This relation was then used to calculate $v_{tb}$, $v_f$ from $v_{tb}^{CM}$, $v_f^{CM}$ for chronic temperature samples that were only registered in the CM frame (Fig. 2, Fig. 5 of the main text). We find that after the applied CM correction (Eqs. 13-15), the newly formed segment boundaries are horizontal in the kymograph (parallel to the anterior edge), as expected for the anterior frame (Fig. A2 B). Additionally, front regression per cycle $\lambda_f = 2\pi v_f/\omega$ in the anterior frame should be approximately equal to the nascent somite size, as confirmed by our measurements (Fig 4. B and D).

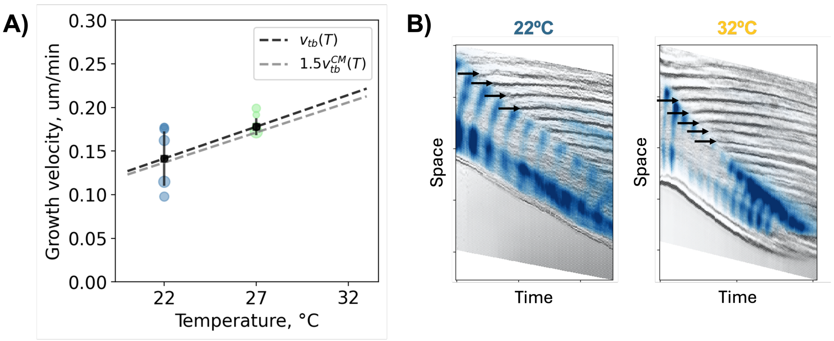


**Figure A2. Measuring the velocity conversion between CM frame and anterior (lab) frame. A)** Elongation velocity measurements in anterior frame (scatter points, linear fit in black) compared to scaled measurements in CM frame (linear fit in grey, same as in Fig. 4B). **B)** Typical examples of kymographs registered in CM frame and converted to anterior frame by adding $v_{CM} = 0.5 v_{tb}^{CM}$. Horizontal arrows indicate nascent somite boundaries.

#### SVD analysis

After posterior registration and cropping, each kymograph $K$ is a rectangular $L*T$ matrix, where the vertical dimension corresponds to space and horizontal to time, so the number of rows $L$ is the length of the ROI in pixels and the number of columns $T$ is the number of time steps. The matrix contains elements $K(x, y)$, $x = 1, ..., L; t = 1, ..., T$. Practically, we have $T < L$.

Singular value decomposition (SVD) factorizes the kymograph $K$ into three matrices: $U$, $\Sigma$, and $V^T$, where $U$ and $T$ are orthogonal matrices and $\Sigma$ is diagonal. For our application of the SVD, a reduced version is sufficient, meaning that we can remove $L - T$ columns from $U$. In the result, $U$ becomes an $L * T$ matrix, while $\Sigma$ and $V$ are square with dimensions $T * T$ (Fig. A3 top).

The SVD effectively provides a representation of $K$ as a sum of rank-1 matrices $K_i$, which we will refer to as modes. The number of these modes is $n = \min(L, T) = T$. Each rank-1 matrix has a very simple structure (Fig. A3): it is an outer product of one column vector from $U$ (spatial component) of length $L$ and one row vector from $V^T$ (temporal component) of length $T$, weighted by the corresponding element of $\Sigma$ - the singular value:

$$K(x, y) = \sum_{i=1}^{n} K_i(x, y) = \sum_{i=1}^{n} s_i\, u_i(x) v_i(t), \tag{16}$$

where $u_i$ is the spatial mode component, $v_i$ is the temporal component, and $s_i$ is the singular value. As the vector components are normalized, the singular value provides the amplitude of each mode, and the modes are always arranged in descending order, $s_{i+1} < s_i$.

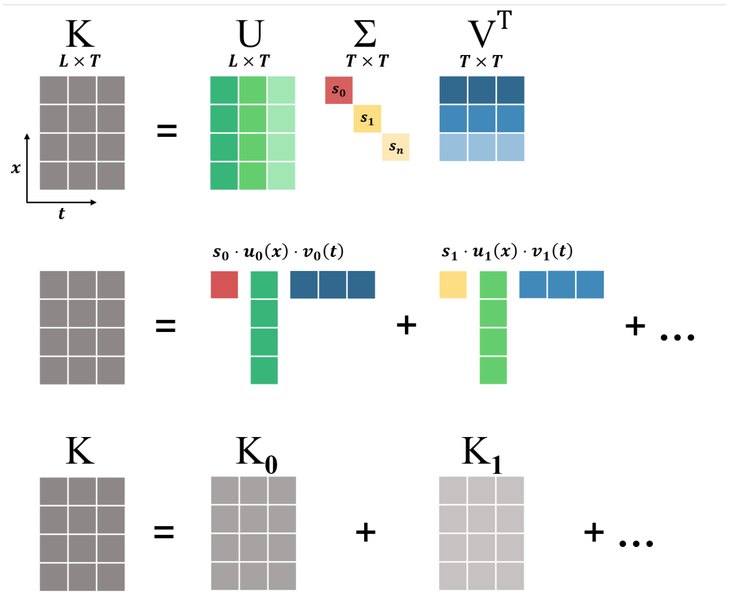


**Figure A3. Singular Value Decomposition**. A kymograph K is decomposed as a product of three matrices: $U$ (spatial mode components), $\Sigma$ (diagonal matrix of singular values), and $V^T$ (temporal mode components). The original $K$ is then a sum of rank-1 matrices (modes) $K_0$, $K_1$, ..., each mode a product of one column of $U$ and one row of $V^T$, weighted by the singular value.

The idea is then to reduce the decomposition based on the mode weights. If a few singular values are much larger than others, we can truncate the sum in Eq. 16. Applying the SVD to our kymographs, we systematically find that $s_1$ is of order $10^3$, $s_2/s_1 \sim 10^{-1}$, and $s_3/s_1 \sim 10^{-2}$ (Fig. 3 D). We therefore approximate the kymograph two SVD modes and set $s_i \to 0$ for $i \geq 3$: $K \approx K_1 + K_2$. Next, we take a closer look at the spatial and temporal components of the obtained modes.

#### Mode fitting

For quantitative analysis, we fit the mode components with functions of space or time as indicated in Table 1. The SVD vectors are determined up to sign, so we distribute the signs between $u_i$ and $v_i$ in a way that the fit parameters are positive.

**Table 1.** Functions used to fit the mode components

| Component | Behavior | Fit |
|---|---|---|
| $u_1(x)$ | Constant | $u_1 = c_1$ |
| $v_1(t)$ | Linear | $v_1 = a_1 t - b_1$ |
| $u_2(x)$ | Nonlinear | $u_2 = a_2/\left(1 + e^{-\beta(x-x_0)}\right) + d_2$ |
| $v_2(t)$ | Constant | $v_2 = -c_2$ |

Combining the components together, we find that the kymograph can be represented simply as

$$K(x,t) \approx const + f(t) + g(x) = const + \omega t - \frac{A}{1 + e^{-\beta(x-x_0)}}. \quad (17)$$

Here we introduce the parameters $\omega = s_1 a_1 c_1$ and $A = s_2 a_2 c_2$, and combine all the constant terms into one. The term $f(t)$ represents a spatially uniform reference oscillator with constant frequency $\omega$. Added to it is a negative spatial phase gradient (phase delay) $g(x)$. In the tailbud, $g(0) \to 0$, so the reference oscillation is directly observed, that is, $\omega$ is the posterior oscillation frequency. The function chosen for $u_2(x)$ is a sigmoid (logistic curve), so the phase gradient $g(x)$ is characterized by the amplitude $A$, steepness $\beta$, and midpoint $x_0$. We have also considered fitting with an exponential function: $u_2(x) = a_2 \exp(\beta x) + d_2$, or $g(x) = -A\left(\exp(\beta x) - 1\right)$. While the two nonlinear functions give rise a largely similar interpretation (see model section below), we differentiate between the two fits by computing the spatial derivative of the phase gradient (Fig. A4). We find that the sigmoid fit and its saturating derivative in the anterior PSM are more consistent with the data.

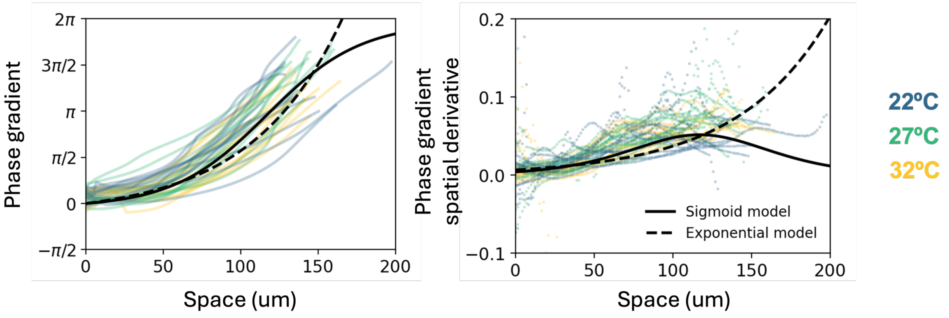


**Figure A4. Choosing between sigmoid and exponential models for the spatial mode fitting Left:** Measured phase gradients $\Delta\phi(x) = s_2\langle v_2(t)\rangle_t u_2(x) + C$. Sigmoid fit (solid): $\Delta\phi = 2\pi/(1+\exp(-0.033(x-117)))$. Exponential fit (dashed): $\Delta\phi = 0.4\exp(0.017x)$. **Right:** Numerical spatial derivatives and the corresponding derivatives of sigmoid and exponential fits.

In fitting nonlinear functions such as $u_2(x)$, the fit parameters are correlated, in particular $\beta$ and $a_2$. We therefore fix $a_2$ for each fitted sample such that the amplitude $A$ is fixed, $A = s_2 a_2 c_2 = 2\pi$. The phase shift of $2\pi$ between anterior and posterior PSM is the maximum observed in medaka. With this constraint, any variability between spatial profiles $g(x)$ in different samples is captured by parameters $\beta$ and $x_0$.

After SVD truncation and fitting, the fit residual of the original matrix $K$ does not exceed 10% and is close to 5% on average. We also check that the fitting does not interfere with the orthonormality property of the SVD component vectors, namely:

$$u_i \cdot u_j = \delta_{ij}, \quad v_i \cdot v_j = \delta_{ij}. \tag{18}$$

For almost all samples, we find that after fitting $u_1 \cdot u_2 \approx 0$, $v_1 \cdot v_2 \approx 0$, and the norm of the fitted components stays close to 1, as expected. Samples with norm values less than 0.9 were excluded from parameter statistics.

Thus, we obtain a parameterization of each kymograph that includes the SVD parameters: segmentation frequency $\omega$, phase gradient exponent $\beta$, as well as velocities: growth velocity $v_{tb}$ and wavefront velocity $v_f$. For analyzing the temperature dependence, we also calculate the following composite parameters: slowdown rate $\alpha$ ($\alpha = \beta v_{tb}$ as defined in the main text), slowdown per cycle $\alpha_c$, regression per cycle $\lambda_f = v_f * \frac{2\pi}{\omega}$, and elongation per cycle $\lambda_{tb} = v_{tb} * \frac{2\pi}{\omega}$.

#### Spatiotemporal modes and the Alpha model

Considering a model of cellular dynamics, we note that the decomposition $\phi(x,t) = f(t) + g(x) +$ const is consistent with a steady-state solution of the Alpha model proposed earlier [15, 22]. In this model, the main components are a reference oscillator $\phi_0$ and a phase gradient $\Delta\phi$. The wave dynamics is driven by the phase difference $\Delta\phi$ between the local phase $\phi(X,t)$ and the reference oscillation in the tailbud $\phi_0(t)$:

$$\phi(X,t) = \phi_0(t) - \Delta\phi(X,t); \tag{19}$$

$$\frac{d\phi_0(t)}{dt} = \omega; \tag{20}$$

$$\frac{\Delta\phi(X,t)}{dt} = F(\Delta\phi). \tag{21}$$

In the original model [22], $F(\Delta\phi) = \alpha\Delta\phi + \varepsilon$, which gives an exponential slowdown of local oscillators with the rate $\alpha$, $\Delta\phi(X,t) \sim \exp(\alpha t)$, and an exponential phase gradient. As the phase gradient is increasing slower than exponentially in the anterior (Fig. A4), we also include a saturating term:

$$\frac{\Delta\phi(X,t)}{dt} = (\alpha\Delta\phi + \varepsilon)\left(1 - \frac{\Delta\phi}{A}\right) \tag{22}$$

The phase shift dynamics can be expressed in terms of a potential $V(\Delta\phi)$:

$$\frac{\Delta\phi(X,t)}{dt} = -\frac{dV(\Delta\phi)}{d\Delta\phi}. \tag{23}$$

A cellular oscillator starts at $\Delta\phi = 0$ (tailbud) and moves in this potential, shifting its phase. For the exponential model, $F(\Delta\phi) \sim \alpha\Delta\phi$, the potential is quadratic, so $\Delta\phi = 0$ is unstable, and any initial phase shift is exponentially amplified with temporal rate $\alpha$. Including the saturation term, $F(\Delta\phi) = \alpha\Delta\phi\left(1 - \frac{\Delta\phi}{A}\right)$, we introduce a stable fixed point at $\Delta\phi = A$ (Fig. A5). We set $A = 2\pi$, a phase shift of one full cycle from the reference (tailbud) oscillator. Furthermore, the phenomenological detuning term $\varepsilon$, $\varepsilon << 1$, provides a small phase shift, effectively tilting the potential around $\Delta\phi = 0$. This ensures that cells exiting the tailbud initially desynchronize from the reference oscillator, $\Delta\phi > 0$.

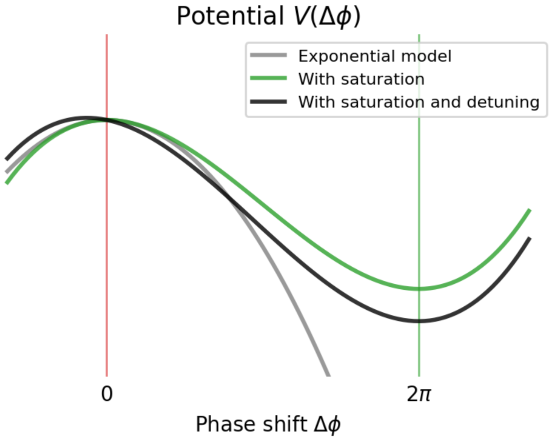


**Figure A5.** Potential for the phase shift dynamics.

For a cell that exits the tailbud at time $t_0$, the resulting phase shift $\Delta\phi(t)$ is

$$\Delta\phi(t) = \frac{A\left(e^{\alpha'(t-t_0)} - 1\right)}{\frac{A\alpha}{\varepsilon} + e^{\alpha'(t-t_0)}}. \tag{24}$$

For the full PSM, Eq. 22 is solved with the moving boundary condition $\Delta\phi(X_{tb}, t) = 0$, $X_{tb}(t) = -v_{tb}t$. Equivalently, new cells are being added in the posterior with $t_0(X) = -X/v_{tb}$, $\Delta\phi(X, t_0(X)) = 0$. The solution is then

$$\Delta\phi(X,t) = \frac{A\left(e^{\alpha'(t+X/v_{tb})} - 1\right)}{\frac{A\alpha}{\varepsilon} + e^{\alpha'(t+X/v_{tb})}} = \frac{A\left(e^{\beta' x} - 1\right)}{\frac{A\alpha}{\varepsilon} + e^{\beta' x}}, \tag{25}$$

where $\alpha' = \alpha + \frac{\varepsilon}{A} \approx \alpha$ and $\beta' = \alpha'/v_{tb} \approx \alpha/v_{tb}$, and we use the distance from the tailbud (PSM coordinate) $x = X + v_{tb}t$.

The resulting phase gradient has a sigmoidal shape:

$$\Delta\phi(x,t) = \frac{A\left(1 - e^{-\beta' x}\right)}{1 + \frac{A\alpha}{\varepsilon}e^{-\beta' x}} \approx \frac{A}{1 + e^{-\beta'(x-x_0)}}, \tag{26}$$

where we define the midpoint

$$x_0 = \frac{1}{\beta'}\log\frac{A\alpha}{\varepsilon}. \tag{27}$$

The analytical solution $\Delta\phi(x,t)$ is therefore the fitted SVD mode (Eq. 17) plus a small exponential correction in the posterior. This correction ensures that the phase shift is strictly $0$ in the tailbud $x = 0$, while for the logistic curve $\Delta\phi \to 0$ as $(x - x_0) \to -\infty$.

The shape of the phase gradient can be characterized by several regions:

- tailbud region - linear build up: $\Delta\phi \approx \frac{\varepsilon}{v_{tb}}x$;
- posterior region - exponential increase: $\Delta\phi \approx \frac{\varepsilon}{\alpha}\left(e^{\beta' x} - 1\right)$;
- midpoint region (with amplitude $A = 2\pi$) - linear gradient: $\Delta\phi \approx \pi + \frac{\pi}{2}\beta'(x - x_0)$.

The spatial exponent $\beta \approx \frac{\alpha}{v_{tb}}$ therefore characterizes the slope of the emergent phase gradient.

The final component of our model is the Her-7 front (wavefront) definition. The Alpha model assumes that the clock and the wavefront are regulated by the same process, that is, the wavefront is also controlled by the phase difference $\Delta\phi$. The wavefront location $X_f$ is given by a specific phase difference $\Delta\phi_*$ to the reference:

$$X_f(t) : \Delta\phi(X_f(t), t) = \Delta\phi_* \tag{28}$$

We find that the PSM length decreases over time ($v_f > v_{tb}$), while the spatial profile $\Delta\phi(x)$ remains constant. This is captured by a simple temporal dependence of the front phase $\Delta\phi_*(t)$ ('cropping' the phase gradient):

$$\Delta\phi(X_f(t), t) = \Delta\phi_*(t) = \Delta_0 - \kappa t. \tag{29}$$

Given a phase gradient $\Delta\phi(X,t)$ and the elongation velocity $v_{tb}$, the front regression velocity is then:

$$v_f(t) = \left|\frac{dX_f(t)}{dt}\right| = \left|\frac{\frac{\partial\Delta\phi(X,t)}{\partial t}}{\frac{\partial\Delta\phi(X,t)}{\partial X}}\right|_{X=X_f(t)} = v_{tb} + \frac{\kappa}{\Phi'(x)}\bigg|_{x=x_f(t)}, \tag{30}$$

where we write the phase shift profile as a moving waveform in the PSM coordinate, $\Delta\phi(X,t) = \Phi(x)$. The front regression is therefore faster than axis elongation, $v_f > v_{tb}$, and $v_f$ depends on the phase profile through its derivative $\Phi'(x)$. The derivative $\Phi'$ increases as the phase gradient is cropped until its midpoint (see Fig. A4), which means $v_f$ and $\lambda_f$ decrease over time, consistent with our experimental observations (Fig. S7 C). In particular, around the midpoint of the gradient, $\Delta\phi(X_f, t) \approx \pi$, we have

$$v_f = v_{tb} + \frac{2\kappa}{\beta\pi}. \tag{31}$$

Then, the front regression per cycle, $\lambda_f = 2\pi v_f/\omega$, is

$$\lambda_f = \lambda_{tb} + \frac{4}{\beta}\frac{\kappa}{\omega}. \tag{32}$$

This means that as the elongation per cycle $\lambda_{tb}$ decreases (i.e. at 32°C), the phase gradient cropping per cycle $\kappa/\omega$ increases to produce a temperature-invariant segment size $\lambda_f$.

#### Model of 6h cycling experiments

In 6h cycling experiments, we observed an increased (average) elongation per cycle. To model the effect of such enhanced elongation, we simulate the PSM as a line of cells where each cellular oscillator (position $X$) evolves according to Eqs. 19-22. At time $t_c$, corresponding to the start of temperature cycles, the elongation velocity is increased by $1.2$. We also hypothesize that the slowdown rate $\alpha$ increases posteriorly to scale with $v_{tb}$, consistent with our observation for chronic temperatures (Fig. 5). We show below that the effects of such parameter changes in the model are: an anterior shift of the phase gradient, a transient increase of PSM length, and a delayed increase of segment size (Fig. A6).

We denote the initial elongation speed $v_0$, and the increased elongation speed $v_1 = v_0 + \Delta v$. At $t = t_c$, the phase gradient is given by the steady state profile:

$$\Delta\phi(X,t) \approx \frac{A}{1 + e^{-\beta(X + v_0 t - x_0)}}, \tag{33}$$

where we use the distance from the tailbud $x = X + v_0 t$. In the control simulation (with constant parameters), this solution remains valid at all times; we denote it as $\Delta\phi^s(X,t)$. In the increased elongation scenario, consider the tailbud coordinate at time $t_c$, $X_{tb}(t_c) = X_c$. For $X > X_c$ (anterior region), $\alpha$ is kept constant, while for $X < X_c$, $\alpha$ is increased: $\alpha_1 = \frac{v_1}{v_0}\alpha_0$. For the simulation in Fig. 6 of the main text, $\alpha = \alpha(X)$ was smoothly interpolated between the two values. For the analytic calculation, we use a step-like change.

For the anterior portion, $X > X_c$, the local oscillators continue to progress with the same $\alpha$, so the solution in the lab frame stays the same: $\Delta\phi(X,t) = \Delta\phi^s(X,t)$, $X > X_c$. We express this solution in the posterior frame using the new PSM coordinate (distance from the tailbud) $x$:

$$x = X - X_c + v_1(t - t_c), \quad t \geq t_c \tag{34}$$

We then have $X + v_0 t - x_0 = x - x_0 - \Delta v(t - t_c) = x - x_0'(t)$, where we define the new midpoint $x_0'(t) = x_0 + \Delta v(t - t_c) > x_0$. Thus, the phase gradient in the anterior region is shifted along the PSM with preserved shape (slope).

In the posterior region, $X < X_c$, the spatial exponent is $\beta_1 = \alpha_1/v_1 = \alpha_0/v_0 = \beta$, so the newly built phase gradient will have the same slope $\beta$ (and approximately the same midpoint $x_0$ since it depends only logarithmically on $\alpha$, Eq. 27).

In PSM coordinate, the anterior (shifted) and posterior (rebuilt) portions of the phase gradient profile are separated by a moving boundary: $X = X_c \Rightarrow x = v_1(t - t_c)$. The full solution for $t > t_c$ is then:

$$\Delta\phi(x,t) = \frac{A}{1+e^{-\beta\left(x-x_0'(t)\right)}}, \quad x > v_1(t-t_c); \tag{35}$$

$$\Delta\phi(x,t) = \frac{A}{1+e^{-\beta(x-x_0)}}, \quad x < v_1(t-t_c). \tag{36}$$

For determining the wavefront, we use the same cropping rule as in the chronic simulation, Eq. 29. As the moving level $\Delta\phi_*(t)$ cuts through the phase gradient $\Delta\phi(X,t)$, different behaviors are observed for $X > X_c$ and $X < X_c$. Initially, $v_f$ is unchanged compared to the control ($X_f(t) = X_f^s(t)$, $X_f > X_c$), producing the same segment size, while the PSM length is gradually increasing compared to the control due to increased elongation. When $X_f$ reaches $X = X_c$ (the entire PSM consists of oscillators with the increased $\alpha$ value), the front accelerates, and the segment size increases. At this point, the phase gradient profile, and thus the PSM length, is recovered.

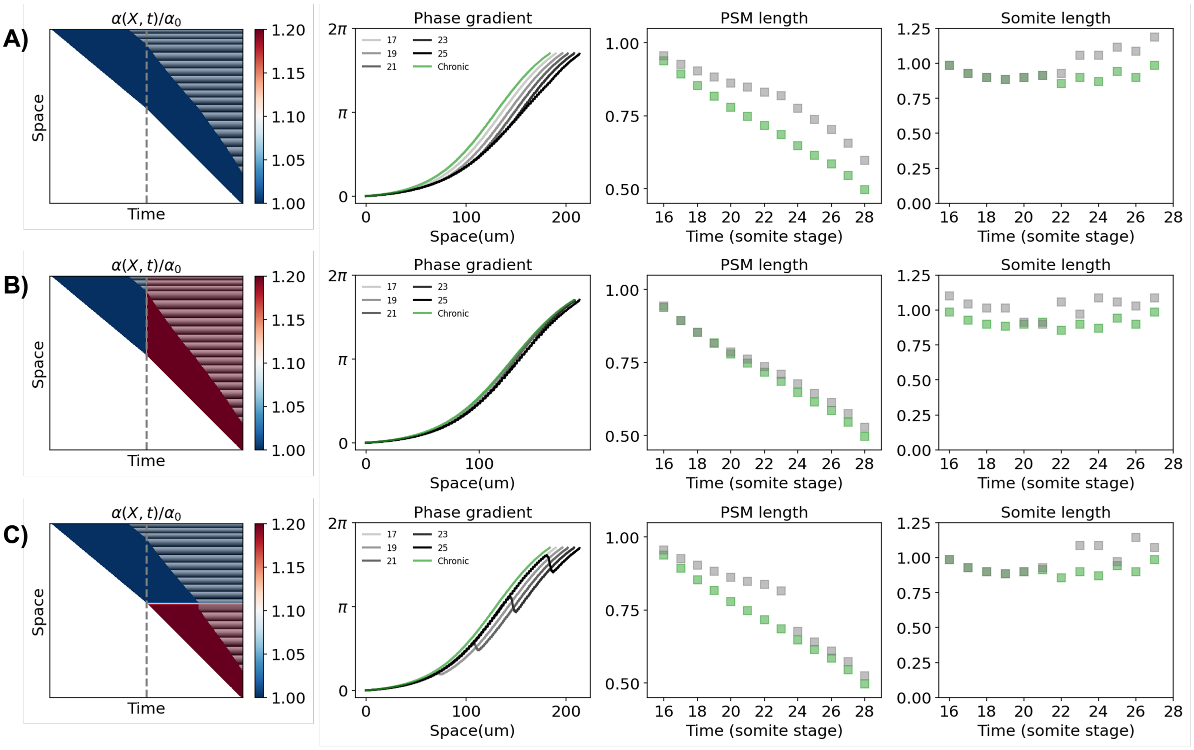


**Figure A6. Simulations of Alpha model with enhanced elongation.** Velocity $v_{tb}$ is increased after the dashed line. The left column shows the colormap of parameter $\alpha$ in the PSM as well as segment boundaries. **A)** No change in $\alpha$: the phase gradient shifts to a lower $\beta$, $\beta_1 = \alpha_0/v_1$. **B)** Global change in $\alpha$: the phase gradient stays constant over time. **C)** Local change of $\alpha$: $\alpha$ is increased only for new posterior cells. The phase gradient is rebuilt with the same $\beta$, Eq. 36.

#### Supplemental Figures

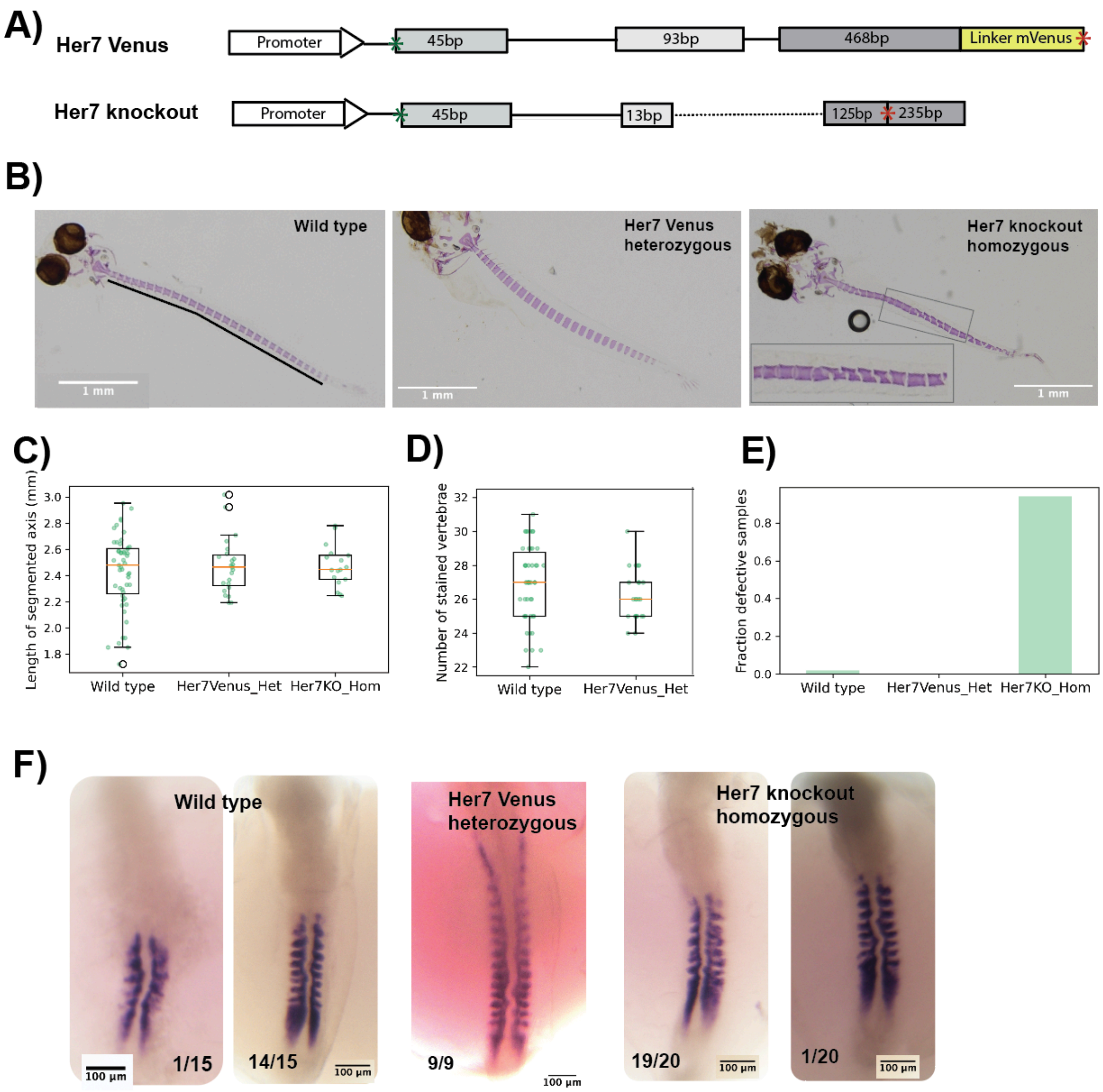


**Figure S1. Validating the Her7 Venus Reporter A)** Schematic showing insertion of mVenus in endogenous Her7 locus in the Her7 Venus genotype, and deletion within endogenous Her7 locus in the Her7 Knockout genotype. The three exons in Her7 locus are color coded by different shades of gray. The dashed line represents the missing region. Green star represents start codon. Red star represents stop codon.. **B)** Representative skeletal staining of hatchlings of the labelled genotypes, raised at 27C. **C)** Length of the segmented axis containing the stained vertebrae in different genotypes (measured as the black line in the first image in B). **D)** Number of stained vertebrae in different genotypes. Due to fusion and irregular shape of vertebrae in Her7 knockout homozygous hatchlings, it is not possible to compute the number of vertebrae accurately. **E)** Fraction of samples having at least one defective vertebrae in each of the three genotypes. N>2, n=26,10,10 for wild type, her7 Venus heterozygous and her7 Knockout homozygous respectively. **F)** Colorimetric in situ hybridization of somite polarity marker (myf5) in embryos of different genotypes. The developmental stages at which embryos were stained are slightly different.

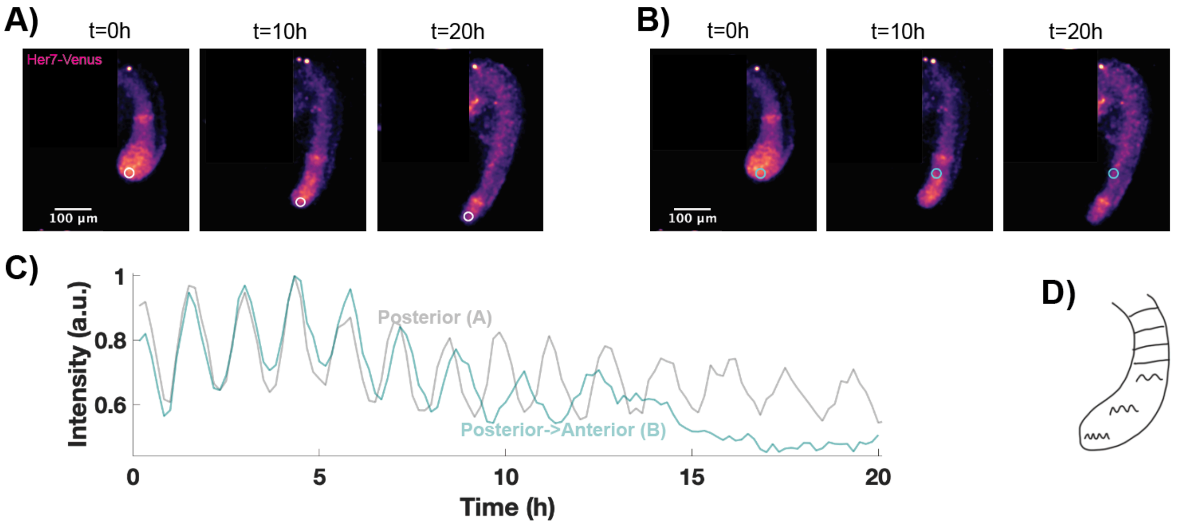


**Figure S2. Slowing down of segmentation clock oscillations in anterior pre-somitic mesoderm A, B)** Her7-Venus heterozygous tail explants during the course of an imaging experiment. **C)** Her7 Venus intensity profiles in the posterior circular region of interest (highlighted in A) and in the circular region of interest that starts in the posterior but gradually becomes more anterior (highlighted in B)). **D)** A schematic showing slowing down of oscillations in the anterior portion of the pre-somitic mesoderm.

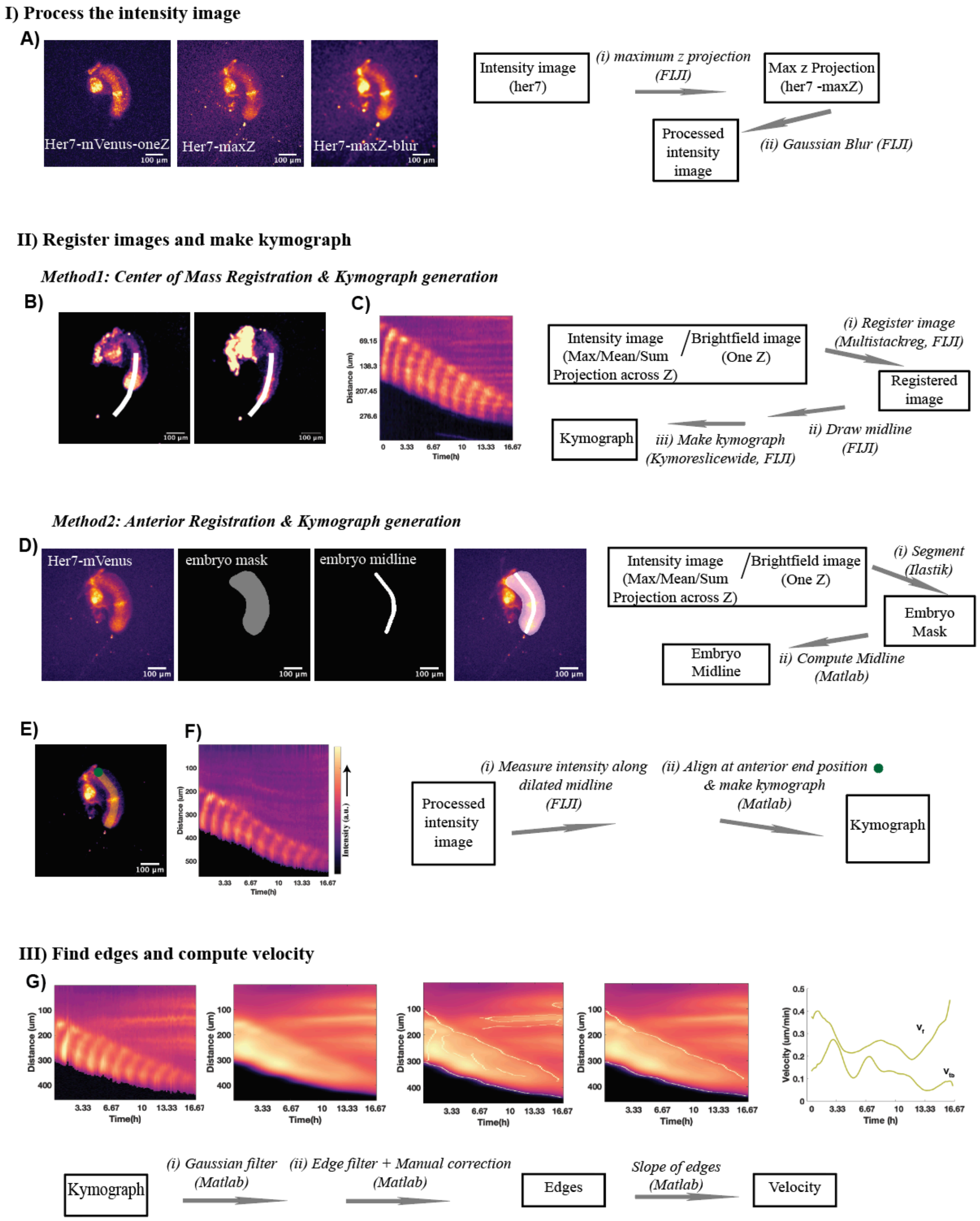
I) Process the intensity image
A)
Her7-mVenus-oneZ
Her7-maxZ
Her7-maxZ-blur
100 µm
Intensity image (her7)
(i) maximum z projection (FIJI)
Max z Projection (her7 -maxZ)
(ii) Gaussian Blur (FIJI)
Processed intensity image
II) Register images and make kymograph
Method1: Center of Mass Registration & Kymograph generation
B)
C)
Distance (um)
Time(h)
Intensity image (Max/Mean/Sum Projection across Z) / Brightfield image (One Z)
(i) Register image (Multistackreg, FIJI)
Registered image
ii) Draw midline (FIJI)
iii) Make kymograph (Kymoreslicewide, FIJI)
Kymograph
Method2: Anterior Registration & Kymograph generation
D)
Her7-mVenus
embryo mask
embryo midline
Intensity image (Max/Mean/Sum Projection across Z) / Brightfield image (One Z)
(i) Segment (Ilastik)
Embryo Mask
ii) Compute Midline (Matlab)
Embryo Midline
E)
F)
Intensity (a.u.)
(i) Measure intensity along dilated midline (FIJI)
(ii) Align at anterior end position & make kymograph (Matlab)
Processed intensity image
Kymograph
III) Find edges and compute velocity
G)
Distance (um)
Time(h)
Velocity (um/min)
Time (h)
Kymograph
(i) Gaussian filter (Matlab)
(ii) Edge filter + Manual correction (Matlab)
Edges
Slope of edges (Matlab)
Velocity

**Figure S3. Representing Her7 oscillation dynamics in the form of intensity kymographs and computing elongation velocity. I)** (A) Raw intensity images along the z-axis are max-projected to obtain one intensity image per time point. **II)** For each sample, images across time are registered, and kymographs are generated using one of two methods: center-of-mass registration or anterior registration. (B, C) Center-of-mass registration: Images are registered using FIJI's MultiStackReg plugin with the center-of-mass registration option. Following registration, a midline is drawn over the oscillating region, and kymographs are subsequently generated using the KymoResliceWide plugin in FIJI. (D–F) Anterior registration: Alternatively, images can be registered using custom-written software that segments intensity images and generates midlines. The midline is then dilated, aligned to the anterior cut site, and used to create intensity kymographs (see Methods). **III)** (G) The edges of the kymograph can then be used to compute elongation velocity (posterior edge) and Her7 front velocity (anterior edge). Under chronic temperature conditions, edges were smoothed and fitted with a straight line to obtain the velocities.

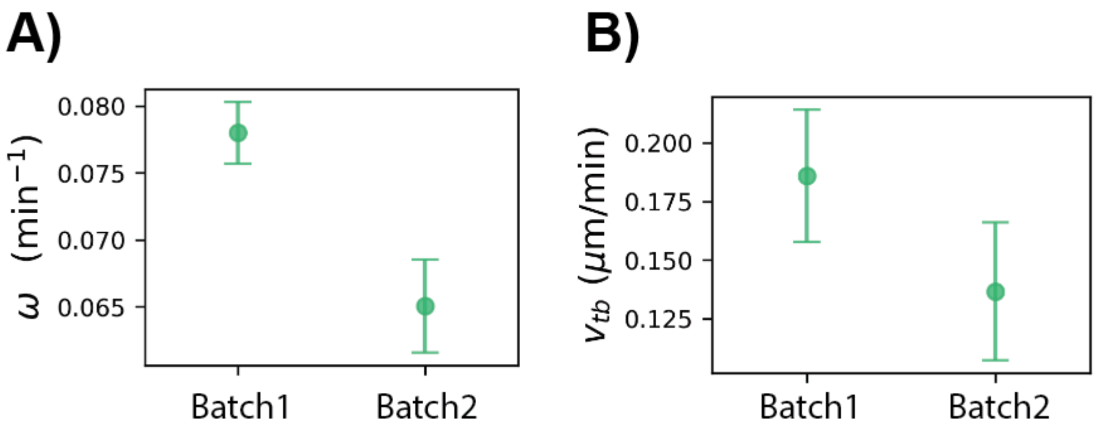


**Figure S4. Difference between the two batches of experiments A, B)** Mean and standard deviation of Segmentation clock frequency $\omega_0$ (A) and Elongation velocity $v_{tb}$ (B) at $27\,^{\circ}\mathrm{C}$ in different batches of experiments (see Methods). For both batches, $N > 3$, $n > 9$.

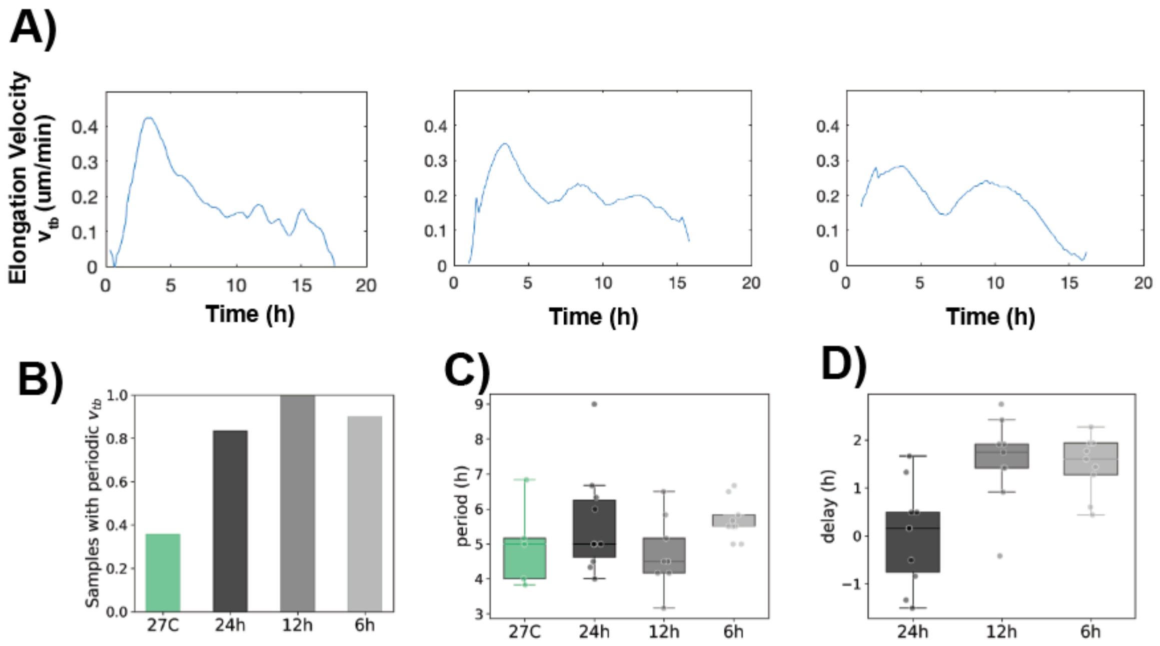


**Figure S5. Elongation velocity in different temperature conditions A)** Three representative profiles of elongation vlocity $v_{tb}$ observed in all temperature conditions. The first profile is classified as non-periodic while the other two as periodic. **B)** Fraction of samples exhibiting periodic $v_{tb}$ in different temperature conditions. $N > 1$, $n > 9$ for all temperatures. **C)** Period of $v_{tb}$, computed as the distance between the first two peaks. This is plotted only for samples with a periodic $v_{tb}$. **D)** Delay, computed as the time between the first temperature peak and the first velocity peak in cycling temperature conditions.

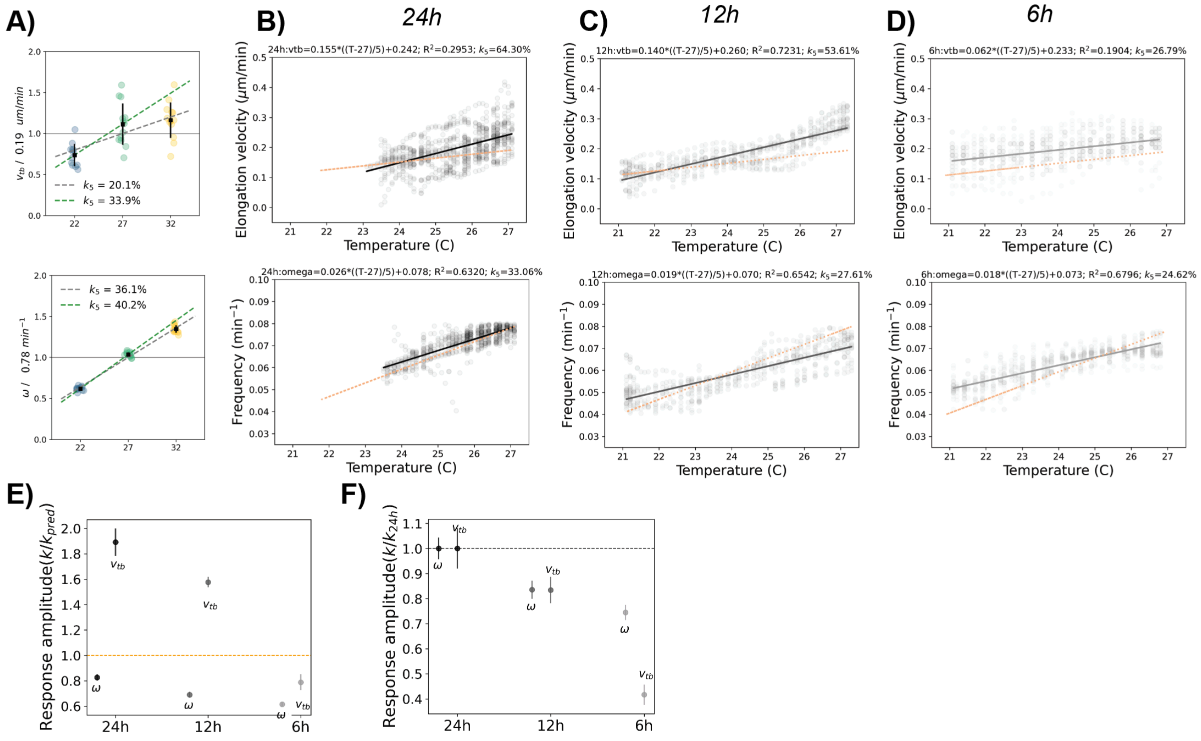


**Figure S6. Amplitude of response of elongation velocity and segmentation clock frequency in cycling temperature conditions A)** Elongation vlocity $v_{tb}$ and segmentation clock frequency $\omega$ in chronic temperature conditions, fitted between 21 and 27C. **B, C, D)** Amplitude of response in $v_{tb}$ and $\omega$ in 24h, 12h and 6h cycling temperature conditions respectively. The orange dotted line represents linear fit for expected values from chronic response. The black solid line represents linear fit of experimental values in the cycling temperatures (listed above the plot). **E, F)** Response amplitude for the $v_{tb}$ and $\omega$, normalized to their response amplitude in expected average chronic temperature (E) and 24h cycling (F) temperatures. For all conditions, $N > 1$, $n > 9$.

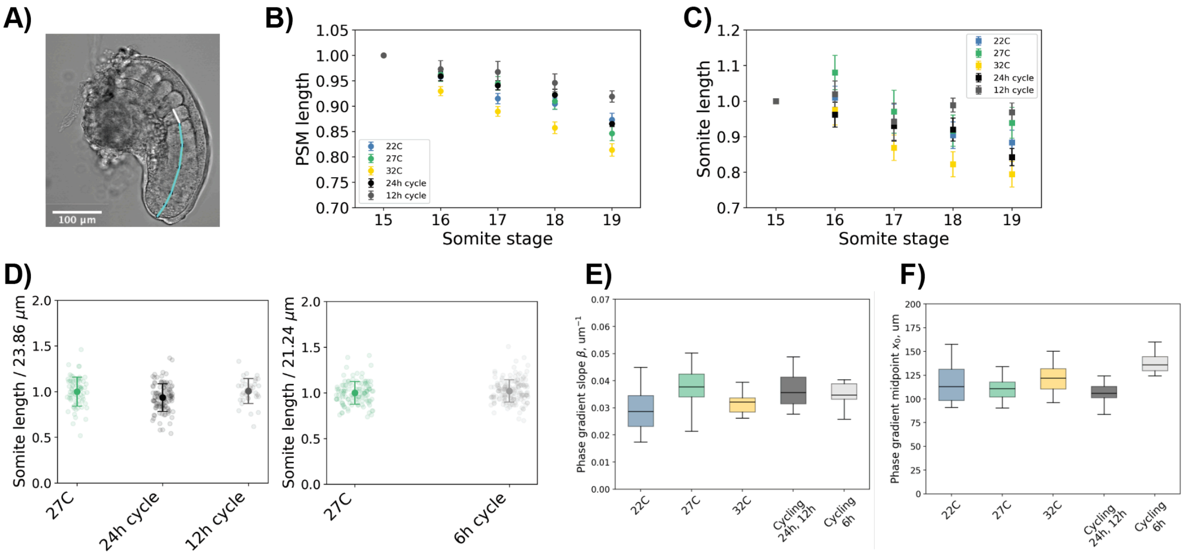


**Figure S7. Tail explant morphology and phase gradient slope measurements in different temperatures A)** A schematic showing measurement of somite length and PSM length in tail explants. White: somite length. Cyan: PSM length. **B, C)** Somite and PSM length progression in individual tail explants in the indicated temperature conditions. The values are normalized to the value at the beginning of the experiment (at somite stage 15) in each tail explant. Error bars represent standard error of mean. **D)** Average somite length during the experiment in different temperature conditions. Experiments are separated by batches. Error bars represent standard error of mean. **E)** Slope of the phase gradient ($\beta$) in different temperature conditions. **F)** Mid point of the phase gradient in different temperature conditions. For all conditions, $N > 2$, $n > 8$.

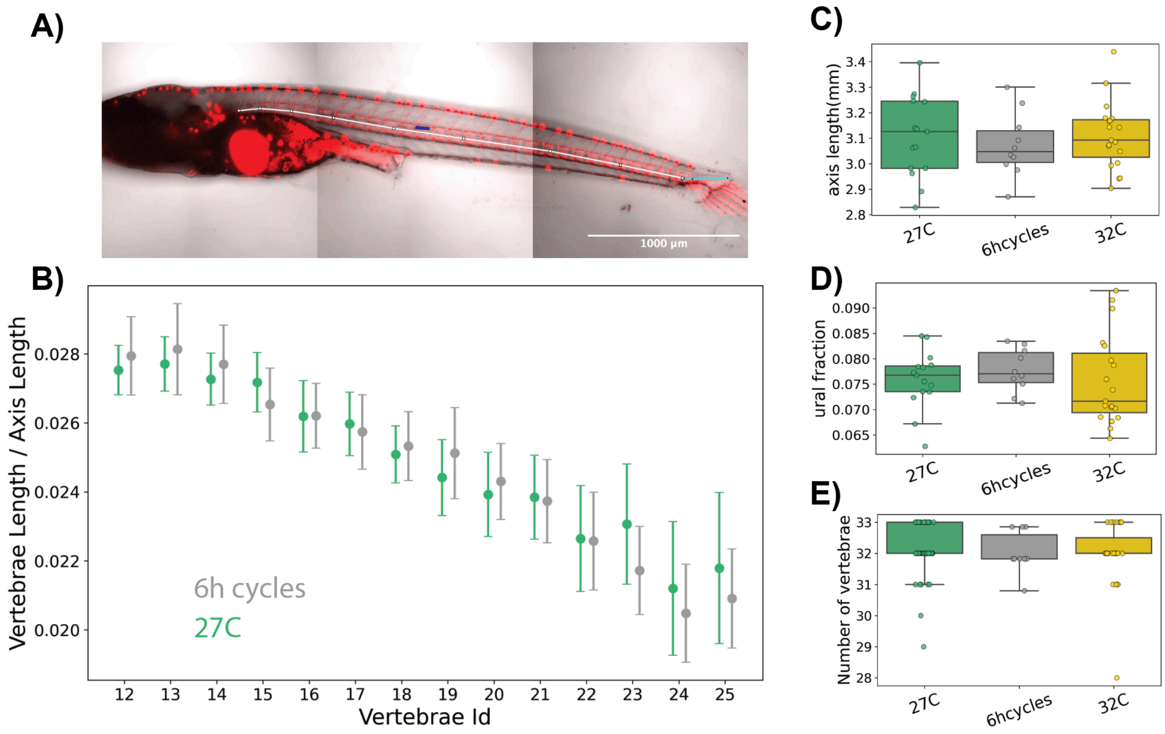


**Figure S8. Hatchling morphology in different temperatures A)** Schematic showing measurement of vertebrae length (blue), axis length (white + cyan) and ural length (cyan) in the hatchlings. **B)** Vertebrae length normalized to axis length in each sample in 27C and 6h temperature conditions. Dots represent mean value across samples, error bars represent standard deviation. Vertebrae Id counted from anterior to posterior. $N = 2, 2$; $n = 9, 10$ for 27C and 6h temperature cycles respectively. **C, D, E)** Box plot showing distribution of axis length, ural fraction and number of vertebrae in the indicated temperature conditions. For C,D: $N = 3, 2, 2$ and $n = 15, 9, 18$ for 27C, 6h cycles and 32C respectively. For E, $N = 2, 2, 3$ and $n = 11, 10, 27$ for 27C, 6h cycles and 32C respectively.
